# Interferon alpha-based combinations suppress SARS-CoV-2 infection in vitro and in vivo

**DOI:** 10.1101/2021.01.05.425331

**Authors:** Aleksandr Ianevski, Rouan Yao, Eva Zusinaite, Laura Sandra Lello, Sainan Wang, Eunji Jo, Jaewon Yang, Erlend Ravlo, Wei Wang, Hilde Lysvand, Kirsti Løseth, Valentyn Oksenych, Tanel Tenson, Marc P. Windisch, Minna Poranen, Anni I. Nieminen, Svein Arne Nordbø, Mona Høysæter Fenstad, Gunnveig Grødeland, Pål Aukrust, Marius Trøseid, Anu Kantele, Astra Vitkauskiene, Nicolas Legrand, Andres Merits, Magnar Bjørås, Denis E. Kainov

## Abstract

There is an urgent need for new antivirals with powerful therapeutic potential and tolerable side effects. In the present study, we found that recombinant human interferon-alpha (IFNa) triggers intrinsic and extrinsic cellular antiviral responses, as well as reduces replication of severe acute respiratory syndrome coronavirus 2 (SARS-CoV-2) in vitro. Although IFNa alone was insufficient to completely abolish SARS-CoV-2 replication, combinations of IFNa with remdesivir or other antiviral agents (EIDD-2801, camostat, cycloheximide, or convalescent serum) showed strong synergy and effectively inhibited SARS-CoV-2 infection in human lung epithelial Calu-3 cells. Furthermore, we showed that the IFNa-remdesivir combination suppressed virus replication in human lung organoids, and that its single prophylactic dose attenuated SARS-CoV-2 infection in lungs of Syrian hamsters. Transcriptome and metabolomic analyses showed that the combination of IFNa-remdesivir suppressed virus-mediated changes in infected cells, although it affected the homeostasis of uninfected cells. We also demonstrated synergistic antiviral activity of IFNa2a-based combinations against other virus infections in vitro. Altogether, our results indicate that IFNa2a-based combination therapies can achieve higher efficacy while requiring lower dosage compared to monotherapies, making them attractive targets for further pre-clinical and clinical development.

## Introduction

Viral diseases continue to pose a serious threat to public health due to a paucity of effective, rapidly deployable, and widely available control measures [1, 2]. Viruses are submicroscopic agents that replicate inside living organisms. When viruses enter and replicate in the cells, viral pathogen-associated molecular patterns (PAMPs) are recognized, and signals are transduced to activate intrinsic and extrinsic immune responses [3]. Pattern recognition receptors (PRRs), including Toll-like receptors (TLRs), RIG-I-like receptors (RLRs) and cytoplasmic DNA receptors sense incoming viruses and activate transcription of IFN genes via NFkB and other pathways. IFNs launch expression of IFN-stimulated genes (ISGs) in infected cells as well as in nearby non-infected cells, protecting them from potential viral invasion. This activation of innate immune response, combined with contributions from adaptive immune response in the host, is often sufficient for elimination of most viruses.

IFNs are a large class of host proteins that are activated during innate antiviral immune response [4, 5]. They are classified into three types, according to the cellular receptor they bind [6] (Fig. S1). Type I IFNs consist of IFN-alpha (IFNa), IFN-beta (IFNb), IFN-epsilon, IFN-kappa and IFN-omega (IFNw) and bind to the IFN-alpha/beta receptor (IFNAR1/2). Type II IFNs consist of IFN-gamma (IFNg) and interact with the IFN-gamma receptor (IFNGR1/2). Finally, type III IFNs, consisting of IFN-lambda-1/IL29 (IFNl1), IFN-lambda-2/IL28A (IFNl2), IFN-lambda-3/IL28B (IFNl3) and IFN-lambda-4 (IFNl4), pass signals through a receptor complex consisting of interleukin 10 receptor 2 (IL10R2) and IFN-lambda receptor (IFNLR1) [7].

Different IFNs induce transcription of different sets of ISGs, which participate in intrinsic antiviral and extrinsic immune responses. For example, ISGs like IFIT1 and OASL activate Ribonuclease L (RNaseL), which leads to the degradation of viral RNA [8]. Moreover, ISGs such as interleukins (ILs), C-X-C and C-C motif chemokines (CXCLs and CCLs) recruit immune cells to the site of infection. Notably, mutations in IFN-signaling pathway genes have resulted in increased susceptibility to viral infections and reduced patient survival [9–12]. However, the exact role of each IFN pathway and their crosstalk remain unclear.

The use of recombinant human IFNs has been approved for treatment of hepatitis C virus (HCV) and hepatitis B virus (HBV) infections [13]. Additionally, IFNs have been shown to be effective against a variety of other viruses in clinical trials and in laboratory settings (Fig. S2) [14–16]. Unfortunately, IFNs possess limited efficacy when administered as antiviral treatments [17, 18] and can cause adverse effects when used at established doses [19].

IFN-related toxicity can be reduced by combining IFNs with other antiviral drugs that act synergistically, thus allowing for the use of smaller doses of each component (Fig. S3). Moreover, synergistic combinations can often have higher efficacy against viral infections than individual components administered as monotherapies, even at lower doses. Indeed, combination treatment of IFNa and ribavirin was the “gold standard” for treatment of chronic HCV infection for more than decade. Similarly, several rhIFN-based drug combinations have been tested against COVID-19. Of note, combinations of IFNb1b/lopinavir–ritonavir/ribavirin, IFNa2b/IFNg, and IFNa/umifenovir were all shown to be effective for treatment of patients with COVID-19 [20–23]. However, despite these promising data, the mode in which IFNs can be optimally combined with other drugs to maximize antiviral and minimize side effects remains unclear.

Here, we have identified several novel synergistic IFNa2a-based drug combinations against SARS-CoV-2, HCV, HEV, FluAV and HIV-1 infections. These treatment combinations are effective at lower concentrations compared to monotherapies. These combinations have powerful treatment potential, which can be leveraged for use in response to imminent viral threats including the emergence and re-emergence of viruses, immune-evading or drug-resistant variants, and viral co-infections.

## Results

### Type I IFNs reduce SARS-CoV-2 replication more efficiently than type II and III IFNs

Although dexamethasone has been shown to improve survival of patients with severe or critical COVID-19 [24], there are currently no curative therapies against SARS-CoV-2. However, previous studies have uncovered several potent antiviral agents, including IFNs, against SARS-CoV-2 *in vitro* and *in vivo* [14, 15, 25]. Here, we tested type I, II, and III IFNs against wild type SARS-CoV-2 (multiplicity of infection (moi) 0.01) in Calu-3 and Vero-E6 cells using cell viability and virus plaque reduction assays as readouts. We observed that type I IFNs rescued both cell types from virus-mediated death and reduced SARS-CoV-2 replication more efficiently than type II and III IFNs. However, the rescue was only partial, and virus replication was reduced only by 2-3 common logarithms (Fig. 1).

**Figure 1.**
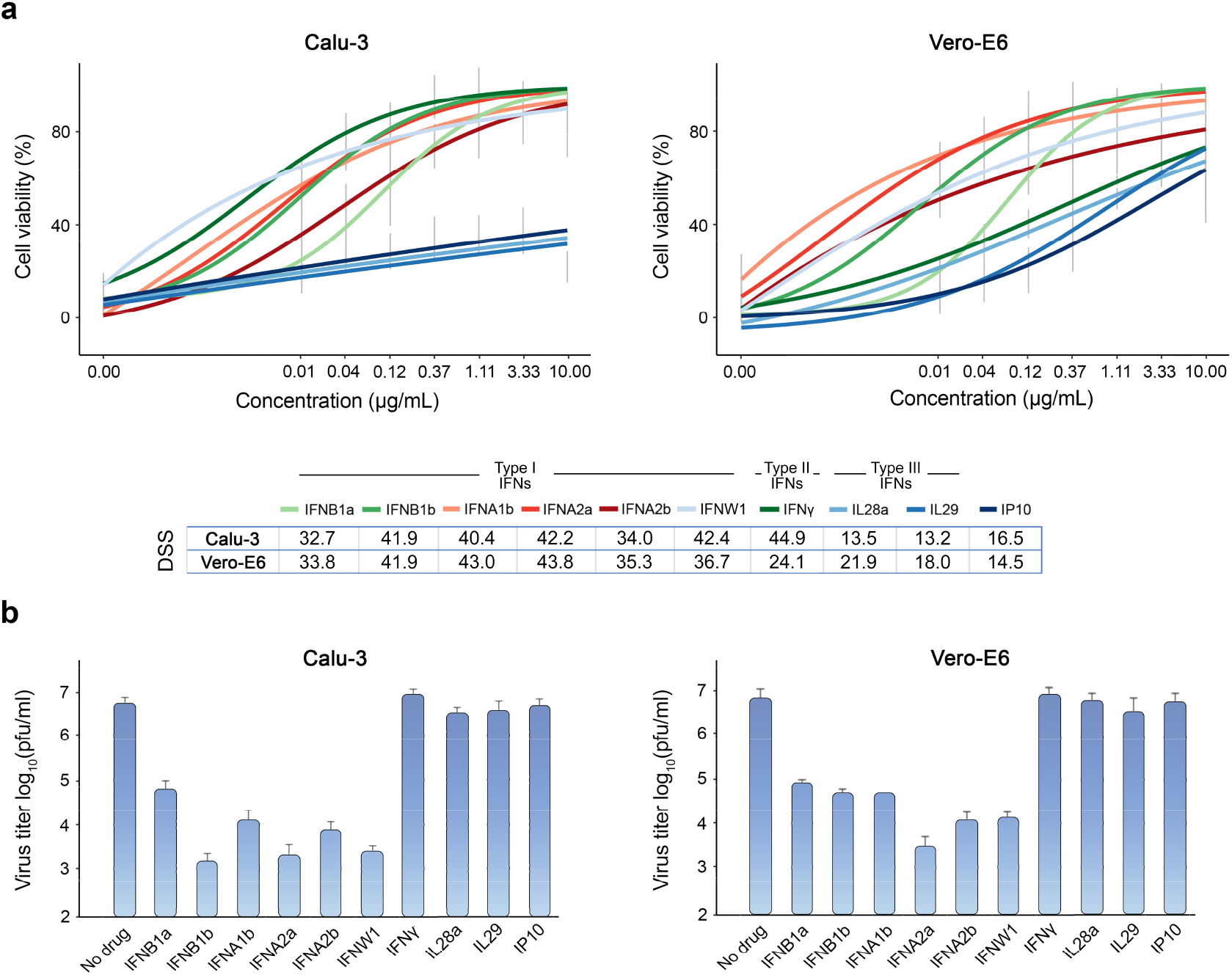
Type I IFNs rescue Calu-3 and Vero-E6 cells from SARS-CoV-2-mediated death and attenuate virus replication. (**a**) The effect of different doses of IFNs on viability of SARS-CoV-2-infected (moi = 0.01) Calu3 and Vero-E6 cells. Cell viability was determined using the CTG assay at 72 hpi. Mean ± SD; n = 3. The anti-SARS-CoV-2 activity of the IFNs was quantified using drug sensitivity scores (DSS). (**b**) The effects of IFNs on viral replication, measured by plaque reduction assay. Mean ± SD; n = 3.

To identify the type I IFN with most activity against SARS-CoV-2 infection, we infected IFN-treated and untreated Calu-3 cells with SARS-CoV-2-mCherry (moi 0.01) and collected media from the cells (p1) after 48 h. The media were diluted 25-fold and applied to noninfected cells for another 48 h (p2). Mock-infected cells were used as controls (Fig. 2a). Fluorescence microscopy, fluorescence intensity analysis, and cell viability assay of p1 and p2 cells showed that IFNa1b, IFNa2a and IFNw1 were more effective inhibitors of SARS-CoV-2 infection than IFNb1a. However, none of the IFNs tested were able to inhibit virus infection completely (Fig. 2b-d).

**Figure 2.**
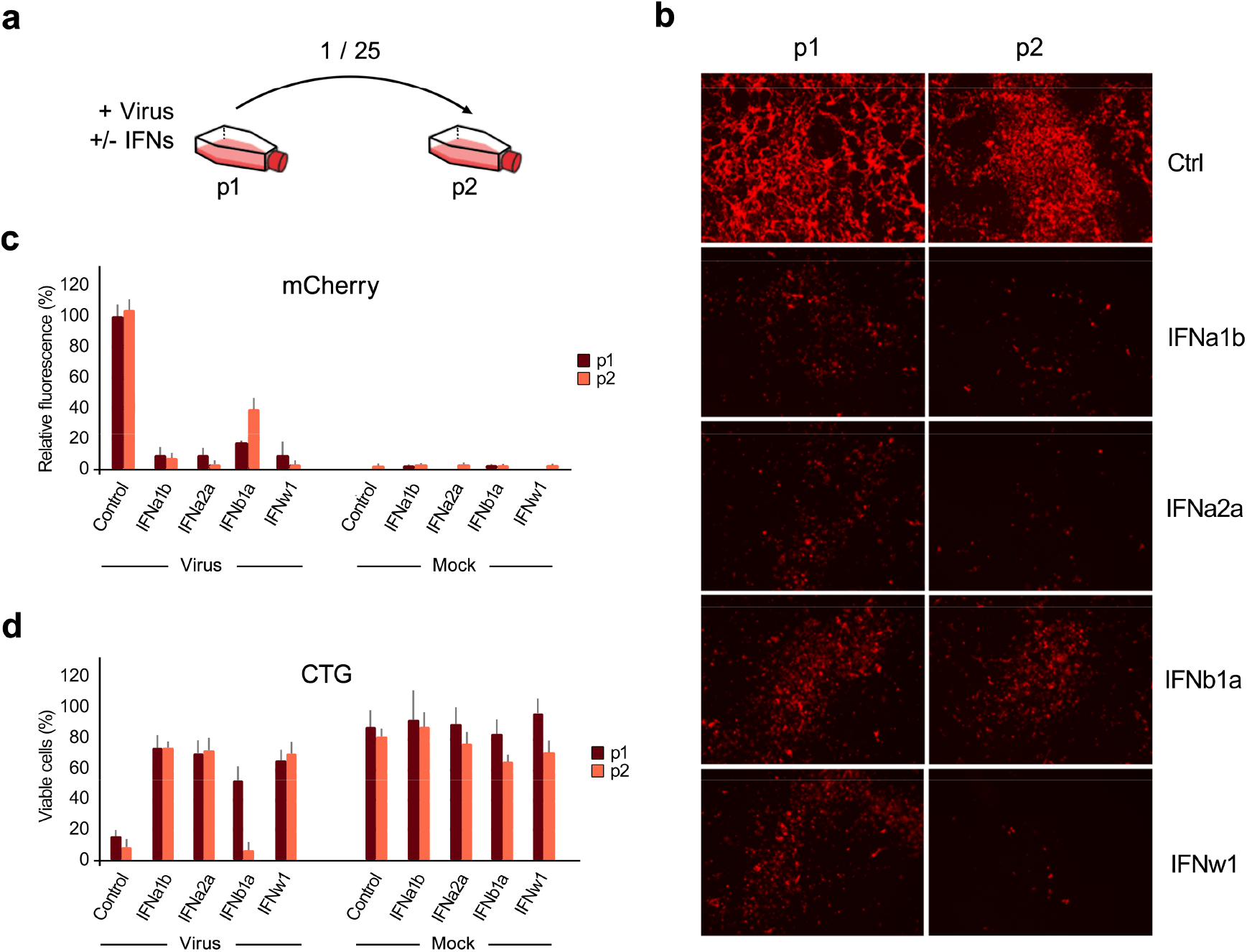
IFNa1b, IFNa2a and IFNw1 are more effective than IFNb1a against SARS-CoV-2-mCherry infection in Calu-3 cells. (**a**) Schematic representation of the experimental setup. (**b**) Fluorescent images of non-treated (Ctrl) and IFN-treated (1 μg/mL) SARS-CoV-2-mCherry-infected Calu-3 cells (p1) and cells (p2) treated with 25-fold diluted media from P1 cells taken at 48 hpi. (**c, d**) Fluorescence intensity and viability analysis of p1 and p2 cells at 48 hpi. Mock-infected cells were used as controls (Mean ± SD; n = 3).

Type I IFNs are encoded by multiple genes and vary slightly from one another in their protein structure. In basic research, IFNa2a is widely used to elucidate the biological activities, structure, and mechanism of action of such type I IFNs. Thus, we next tested IFNa2a against various doses of SARS-CoV-2-mCherry and different time of drug addition. The Calu-3 cells were treated with 1 μg/mL IFNa2a at indicated time points, then infected with SARS-CoV-2-mCherry at indicated moi. After 48 h, fluorescence intensity and cell viability analysis were performed. We found that efficacy of IFNa2a treatment in preventing SARS-CoV-2 infection is dependent on virus load, decreasing in efficacy as moi increases (Fig. 3a) as well as on time of addition, showing more efficacy when given prior virus infection than following infection (Fig. 3b).

**Figure 3.**
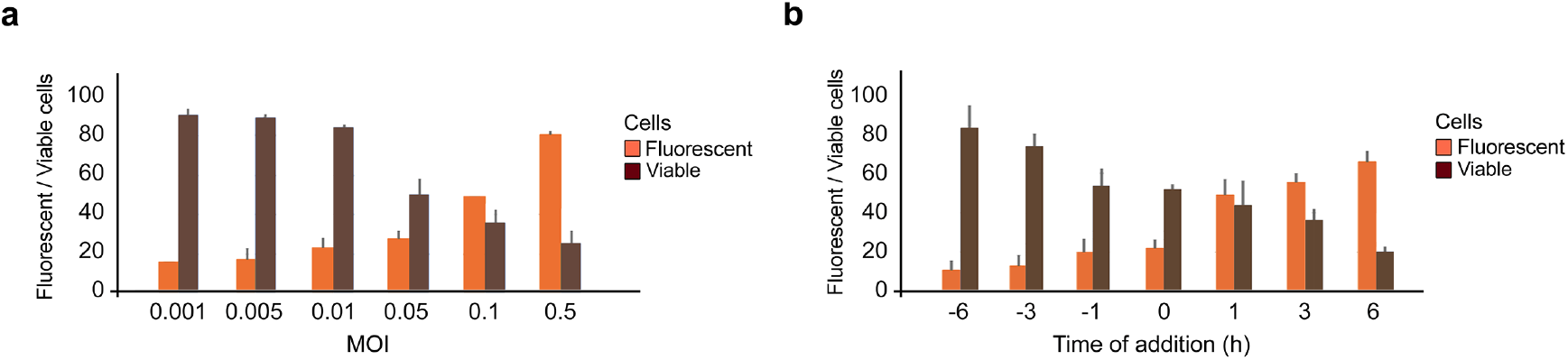
Anti-SARS-CoV-2 activity of IFNa2a depends on moi and time of administration. (**a**) Calu-3 cells were treated with 1 μg/mL IFNa2a and infected with indicated moi of SARS-CoV-2-mCherry. Fluorescence intensity and cell viability were measured after 48 h (Mean ± SD; n = 3). (**b**) Calu-3 cells were treated with 1 μg/mL IFNa2a prior, simultaneously or post infection with SARS-CoV-2-mCherry (moi 0.01). Fluorescence intensity and cell viability were measured after 48 h (Mean ± SD; n = 3).

### IFNa2a reduces the SARS-CoV-2 RNA synthesis and promotes virus-mediated induction of type III IFNs, IFNb1 and ISGs

To shed new light on the mechanism of action of IFNa2a, we evaluated their effect on expression of cellular genes and transcription of viral RNA in mock- and SARS-CoV-2-infected Calu-3 cells. For this, cells were treated with 1 μg/mL of IFNa2a, other type I IFNs, or vehicle; then infected with virus or mock. After 24 h, we analyzed polyadenylated RNA using RNA-sequencing. We found that IFNa2a and other type I IFNs attenuated production of viral RNA (Fig. 4a), while increasing expression of many ISGs in cells, regardless of virus- or mock-infection (Fig. 4b). These ISGs include IFIT1, IFIT2 and IFIT3, which play a role in recognition of viral RNA; OASL and OAS2, which are involved in RNase L-mediated RNA degradation; and IDO1, which is essential for kynurenine biosynthesis [26–29]. Interestingly, IFNa2a and other type I IFNs boosted virus-activated expression of type III IFNs (IFNl1, IFNl2, IFNl3 and IFNl4) as well as IFNb1, which belongs to type I IFN. These results indicate that IFNa2a does not only trigger expression of ISGs, but also amplifies expression of other IFNs usually activated by viral infections, creating a positive feedback loop of IFN signaling during SARS-CoV-2 infection.

**Figure 4.**
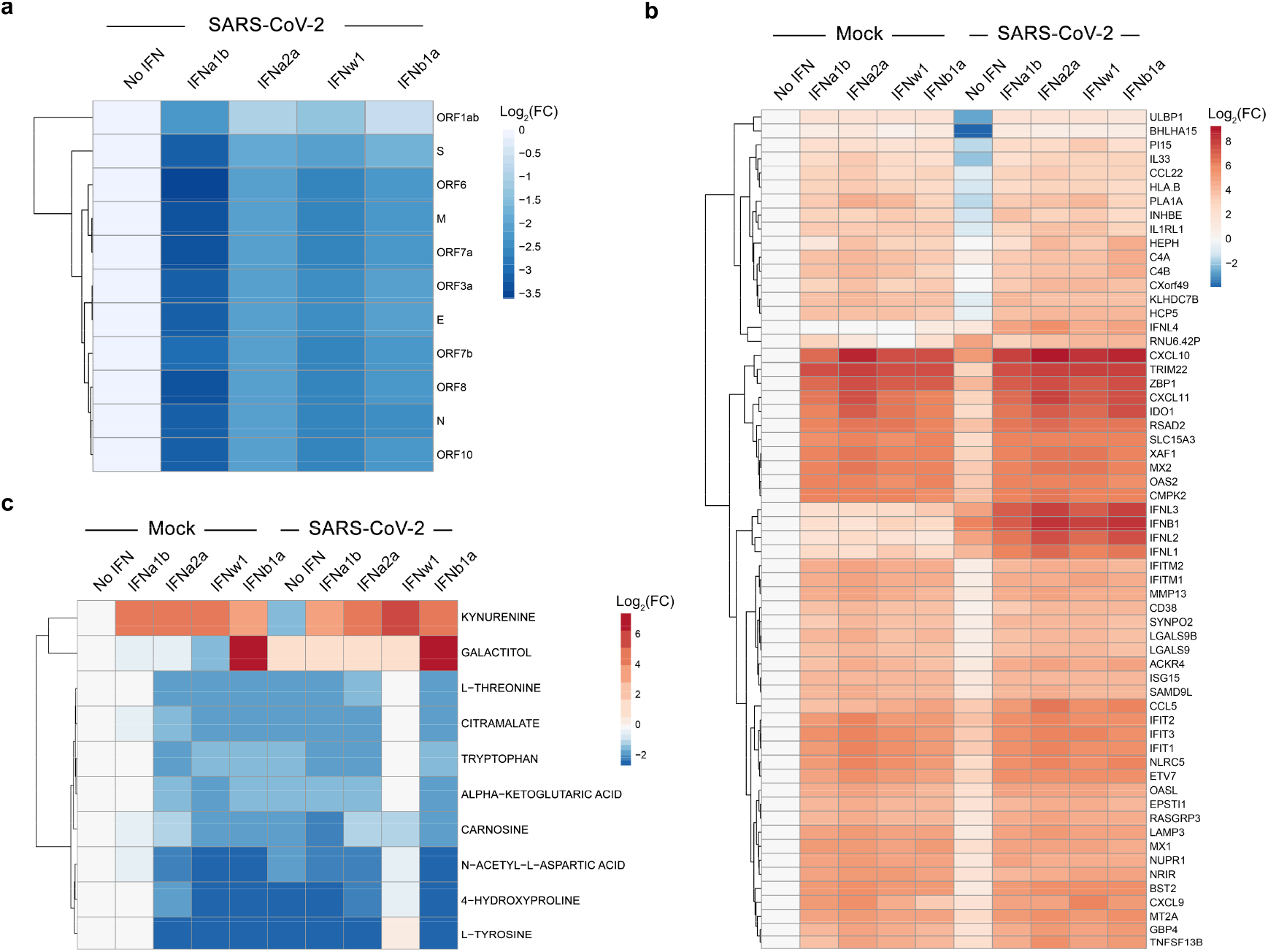
Transcriptomic and metabolomic analysis of mock- and SARS-CoV-2-infected Calu-3 cells non-treated or treated with type I IFNs. (**a**) Calu-3 cells were stimulated with IFNs (1 μg/mL) or non-stimulated and infected with SARS-CoV-2 (moi = 0,01). A heatmap of viral RNAs affected by treatment is shown. Each cell is colored according to the log_2_-transformed expression values of the samples, expressed as fold-change relative to the nontreated control. (**b**) Calu-3 cells were either stimulated with purified recombinant human IFN (1 μg/mL) or left untreated with IFN, then infected with either mock or SARS-CoV-2 (moi = 0,01). A heatmap of the most variable cellular genes affected by treatment and virus infection is shown. Each cell is colored according to the log_2_-transformed expression values of the samples, expressed as fold-change relative to the nontreated mock-infected control. (**c**) Cells were treated as for panel b. After 24 h, the cell culture supernatants were collected, and metabolite levels were determined by LC-MS/MS. A heatmap of the most affected metabolites is shown. Each cell is colored according to the log_2_-transformed profiling values of samples, expressed as fold-change relative to the mock control.

Next, we studied the effect of IFNa2a on the metabolism of mock- and SARS-CoV-2-infected Calu-3 cells. A total of 93 mainly polar metabolites were quantified at 24 hpi (Fig. S4). We found that tyrosine and 4-hydroxyproline levels were substantially lowered during viral infection (log_2_FC<-2). Additionally, administration of IFNa2a or other type I IFNs lowered the levels of several metabolites including tryptophan while increasing kynurenine, regardless of viral infection (log_2_FC>3; Fig. 4c). This indicates that IFNa2a activates IDO1-mediated kynurenine biosynthesis, which could be associated with adverse reactions such as suppression of T-cell responses, pain hypersensitivity and behavior disturbance [30].

### Synergistic IFNa2a-based combinations against SARS-CoV-2 infection in vitro and in vivo

Next, we examined whether IFNa2a in combinations with known SARS-CoV-2 inhibitors remdesivir, EIDD-2801, camostat, cycloheximide, or convalescent serum [31–35] can protect cells from virus infection more efficiently and at lower concentrations than IFNa2a alone. Remdesivir and EIDD-2801 are nucleoside analogues which inhibit viral RNA synthesis [33, 36]. Camostat, a serine protease inhibitor, reduces SARS-CoV-2-cell membrane fusion by binding host TMPRSS2 [37]. In addition, camostat possesses some potential beneficial immunomodulatory effects by interfering with the bradykinin/kallikrein pathways [38]. Cycloheximide inhibits translation elongation and, thereby, reduces SARS-CoV-2 replication [32]. Convalescent serum contains neutralizing antibodies which bind S protein of SARS-CoV-2 preventing virus entry into the cells [16].

We first confirmed antiviral activities of these known viral inhibitors on Calu-3 cells using SARS-CoV-2-mCherry (Fig. 5a, Fig. S5a). Then, we tested the antiviral efficacy and toxicity of these agents in combination with IFNa2a in Calu-3 cells by monitoring virus-mediated mCherry expression and cell viability (CTG). Each drug combination was tested in a 6×6 dose-response matrix, where 5 doses of single drugs are combined in a pairwise manner. As a result, we obtained dose-response matrices demonstrating virus inhibition and cell viability achieved by each combination (Fig 5c,d; Fig. S5b-e). We plotted synergy distribution maps, showing synergy (higher than expected effect) at each pairwise dose. For each drug pair, we calculated average ZIP synergy scores for the whole 6×6 dose-response matrices and for most synergistic 3×3 dose-regions, summarizing combination synergies into single metrics (Fig. 5e). We observed that all combinations showed a strong synergy (synergy scores >10) at various combination doses. Thus, the observed synergy allows us to substantially decrease the concentration of both components to achieve antiviral efficacy that was comparable to those of individual drugs at high concentrations.

**Figure 5.**
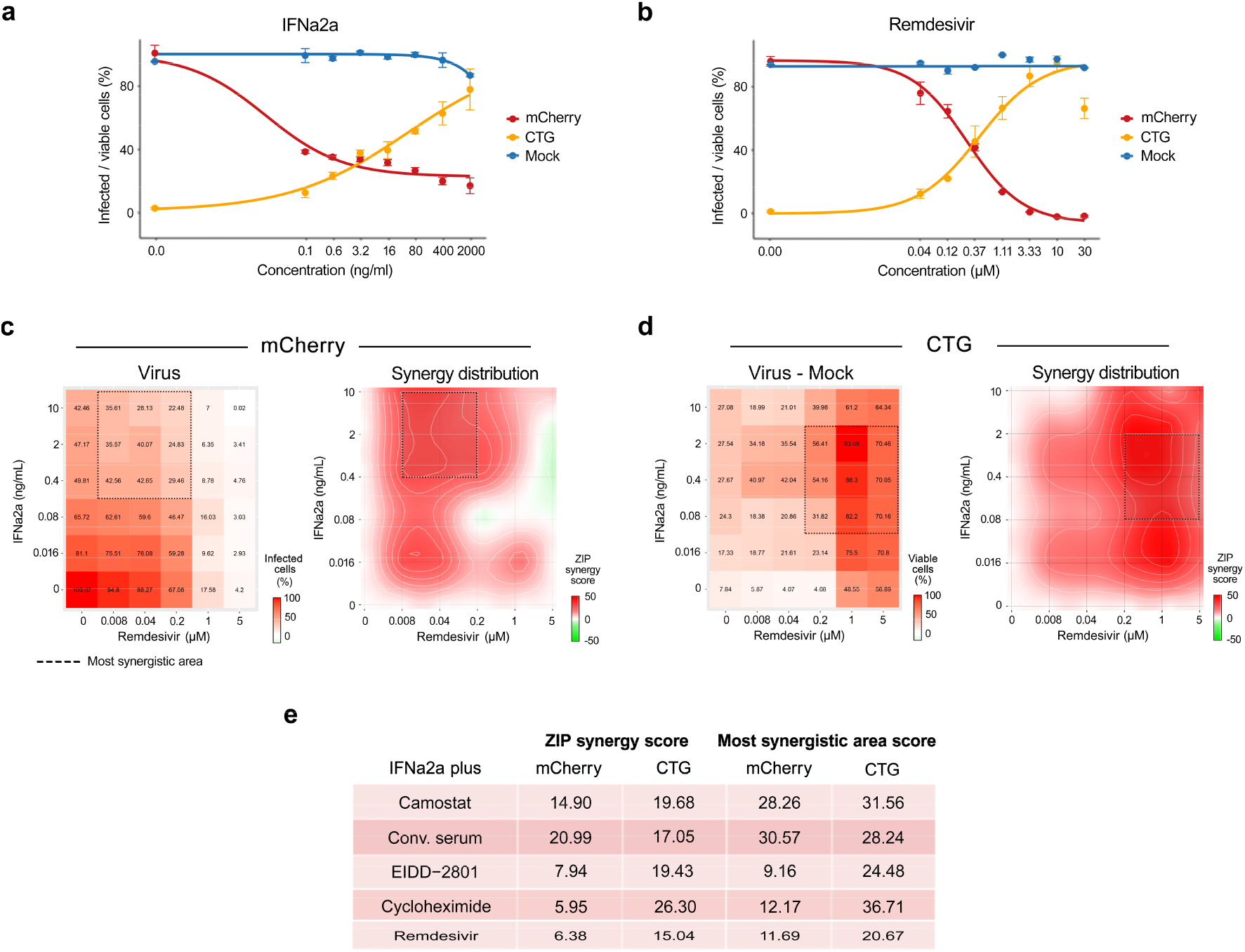
Synergistic IFNa2a-based combinations against SARS-CoV-2-mCherry infection in Calu-3 cells. (**a,b**) Calu-3 cells were treated with increasing concentrations of IFNa2a or remdesivir and infected with the SARS-CoV-2-mCherry or Mock. After 48 h, the virus-mediated mCherry expression was measured (red curves). After 72 h, viability of virus- and mock-infected cells was determined using a CTG assay (yellow and blue curves, respectively). Mean ± SD; n = 3. (**c**) The 6 × 6 dose-response matrices and interaction landscapes of IFNa2a and remdesivir obtained using fluorescence analysis of SARS-CoV-2-mCherry-infected Calu-3 cells. ZIP synergy score was calculated for the drug combinations. (**d**) The 6 × 6 dose-response matrices and interaction landscapes of IFNa2a and remdesivir obtained using a cell viability assay (CTG) on mock-, and SARS-CoV-2-mCherry-infected Calu-3 cells. The selectivity for the indicated drug concentrations was calculated (selectivity = efficacy-(100-Toxicity)). ZIP synergy scores were calculated for indicated drug combinations. (**e**) ZIP synergy scores (synergy score for whole 6×6 dose-response matrices) and the most synergistic area scores (synergy score for most synergistic 3×3 dose-regions) calculated for indicated drug combinations.

Both remdesivir and *rh*IFNa2a (Pegasys) were approved for the treatment of COVID-19 infection in several countries. Therefore, we evaluated the antiviral effect of IFNa2a-remdesivir combination on iPSC-derived lung organoids (LOs). Thirty-day-old LOs were treated with 5 ng/mL IFNa2a, 0.5 μM remdesivir, or a combination thereof, then infected with SARS-CoV-2-mCherry. At 72 hpi, the organoids were analyzed for viral reporter protein expression (mCherry) and cell death (CellToxGreen). We found that IFNa2a-remdesivir substantially attenuated virus-mediated mCherry expression without affecting cell viability (Fig. 6a).

**Figure 6.**
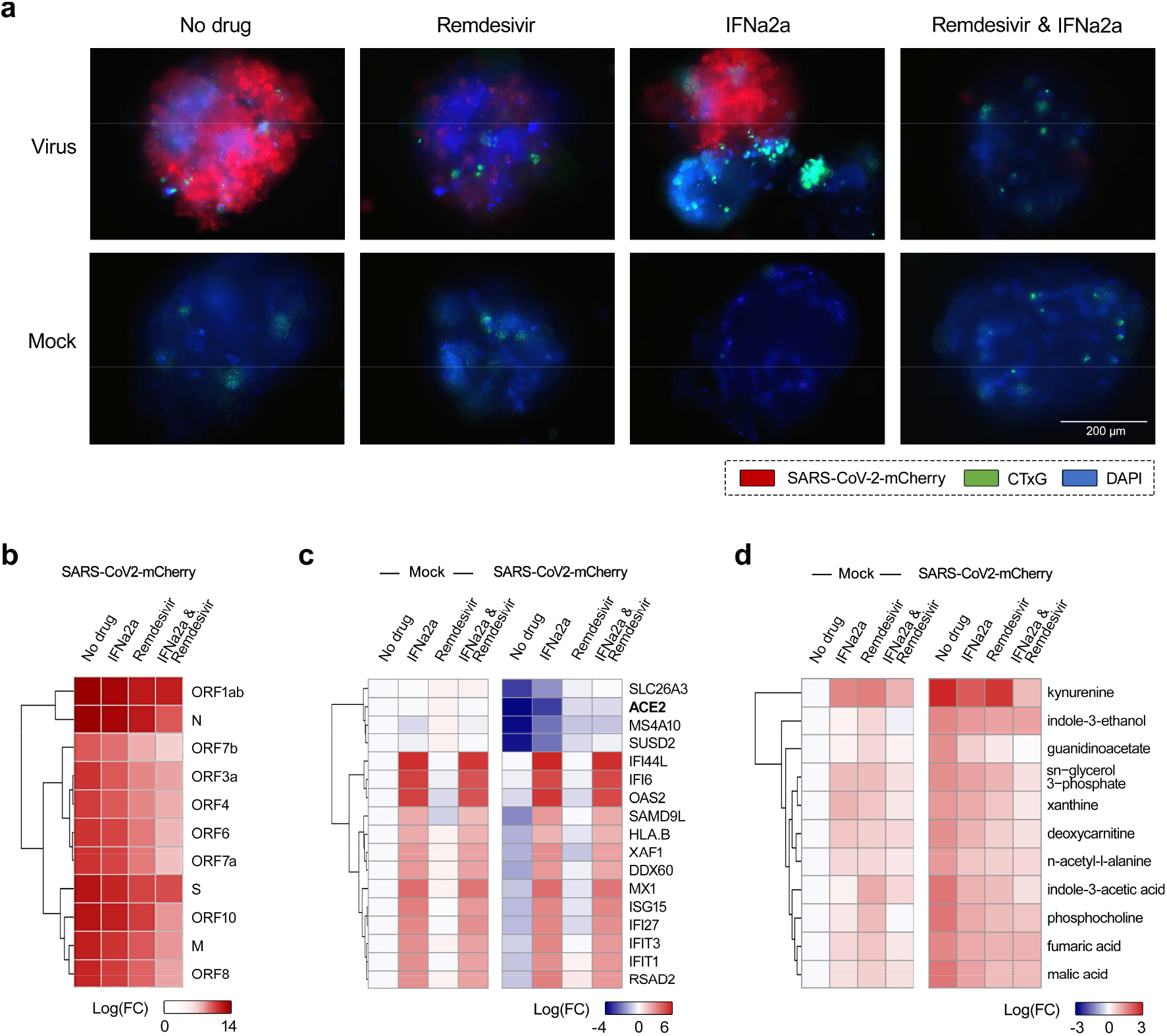
Evaluation of antiviral effect of IFNa2a-remdesivir combination in human lung organoids (LOs). (**a**) LOs were treated with 0,5 μM remdesivir, 5 ng/mL IFNa2a, their combination or vehicle, and infected with SARS-CoV-2-mCherry (moi = 0,1) or mock. Fluorescence of drug- or carrier-treated SARS-CoV-2-mCherry-infected LOs was detected at 48 hpi. Virus infection, cell nuclei, and cytotoxicity are shown in red, blue, and green, respectively. Scale bars, 200 μm. (**b**) LOs were treated with 0,5 μM remdesivir, 5 ng/mL IFNa2a, their combination or vehicle, and infected with SARS-CoV-2-mCherry (moi = 0,1). After 48 h, total RNA was extracted and sequenced. A heatmap of viral RNAs affected by treatment is shown. Each cell is colored according to the log2-transformed expression values of the samples, expressed as log2 fold-change relative to the nontreated control. (**c**) LOs were treated and infected as for panel a. After 48 h, total RNA was extracted and sequenced. A heatmap of the most variable cellular genes affected by treatment and virus infection is shown. Each cell is colored according to the log2-transformed expression values of the samples, expressed as fold-change relative to the nontreated mock-infected control. Cut-off - 3.75. (**d**) Cells were treated as for panel a. After 48 h, the cell culture supernatants were collected, and metabolite levels were determined by LC-MS/MS. A heatmap of the most affected metabolites is shown. Each cell is colored according to the log2-transformed profiling values of samples, expressed as fold-change relative to the mock control. Cut-off - 1.5.

We also evaluated the effect of the combination treatment on viral and cellular RNA expression in LOs. RNA-sequencing revealed that at 48 hpi IFNa2a-remdesivir substantially reduced production of viral RNA by contrast to single agents (Fig. 6b). Treatment with IFNa2a-remdesivir also led to elevated levels of ACE2 and other genes involved in lipid metabolism (APOA4, ADH4, CYP3A7, PON3, FADS6, SDR16C5, ENPP7, FABP2, CUBN, and SERPINA6) [39, 40], for which transcription was substantially down-regulated during SARS-CoV-2 infection (Fig. S6, Fig. 6c). Importantly, the set of IFNa2a-induced ISGs in LOs is consistent to what we observed in Calu-3 cells (Fig. S6).

Furthermore, we studied the effect of IFNa2a-remdesivir on the metabolism of SARS-CoV-2- and mock-infected LOs. A total of 82 metabolites were quantified in LO culture supernatants at 48 hpi. Administration of IFNa2a-remdesivir prevented virus-mediated alteration of metabolism, excluding kynurenine biosynthesis (Fig. S7, Fig. 6d), which is in line with the results obtained on IFNa2a-stimulated Calu-3 cells.

Next, we examined whether IFNa and remdesivir can affect the replication of SARS-CoV-2 *in vivo*. Four groups of 8 six-week-old female Syrian hamsters were injected IP with recombinant mouse IFNa, remdesivir, IFNa-remdesivir combination or vehicle thereof. After 2 h of drug treatment, animals received SARS-CoV-2 intranasally. After 3 days, animals were anesthetized and euthanized, and the lungs were collected (Fig. 7a). Virus titers from hamster lung homogenates in each treatment group were determined using plaque reduction assays (Fig. 7b). In addition, viral RNA was extracted and sequenced. Sequencing results were validated using RT-qPCR (Fig. 7c,d). The IFNa-remdesivir combination attenuated the SARS-CoV-2 production and the synthesis of some viral RNAs more efficiently than individual agents.

**Figure 7.**
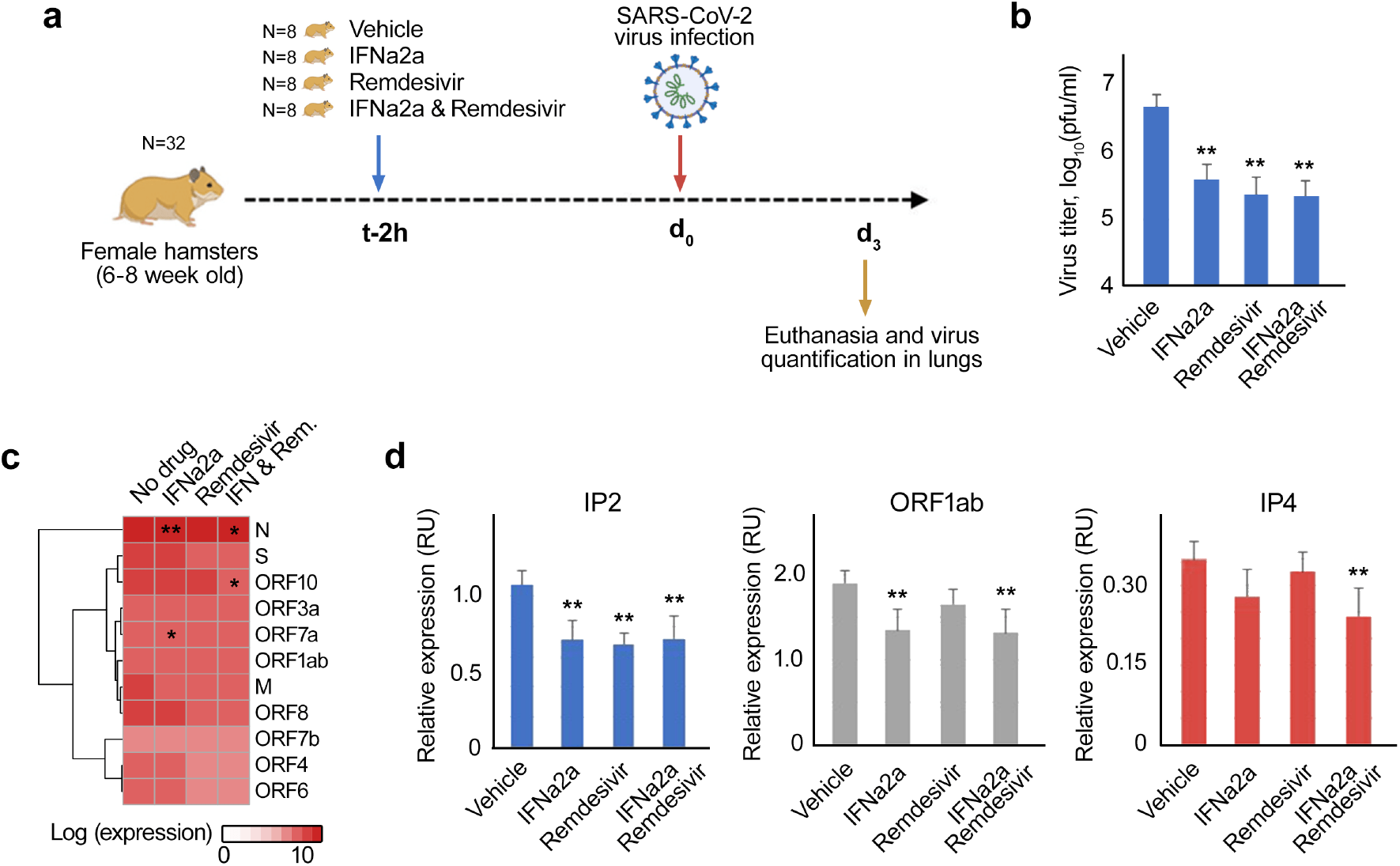
Evaluation of antiviral activity of recombinant mouse IFNa-remdesivir combination in vivo. (**a**) Schematic representation of the experimental setup. (**b**) The effects of IFNa-remdesivir combination on viral replication in hamster lungs, measured by plaque reduction assay. Mean ± SD; n = 8. (**c**) A heatmap of viral RNAs affected by treatment. Each cell is colored according to the log2-transformed expression values of the samples, expressed as log2 fold-change relative to the nontreated control. Mean, n = 8. (**d**) RT-qPCR analysis of selected viral RNA. Expression of viral RNA was normalized to b-actin control. Mean ± SD, n = 8. Statistically significant differences in viral gene expression between non-treated and treated animals are indicated with asterisks (**p<0.05, *p<0.1, Wilcoxon test).

### Synergistic IFNa2a-based combinations against other viral infections

To extend our findings beyond SARS-CoV-2, we tested IFNa2a in combination with known HCV inhibitors, sofosbuvir and telaprevir, using GFP-expressing HCV in infected Huh-7.5 cells. Sofosbuvir is a nucleoside analogue, which inhibits viral RNA synthesis, whereas telaprevir is an orally available peptidomimetic that targets the HCV serine protease and disrupts processing of viral proteins and formation of a viral replication complex. Eight different concentrations of the compounds alone or in combination were added to virus- or mock-infected cells. HCV-mediated GFP expression and cell viability were measured after 72 hpi to determine compound efficacy and toxicity. Both IFNa2a-sofosbuvir and IFNa2a-telaprevir lowered GFP-expression without detectable cytotoxicity at indicated concentrations with synergy scores of 3 and 5 (the most synergistic area scores: 14 and 16), respectively (Fig. S8, Fig. 8).

**Figure 8.**
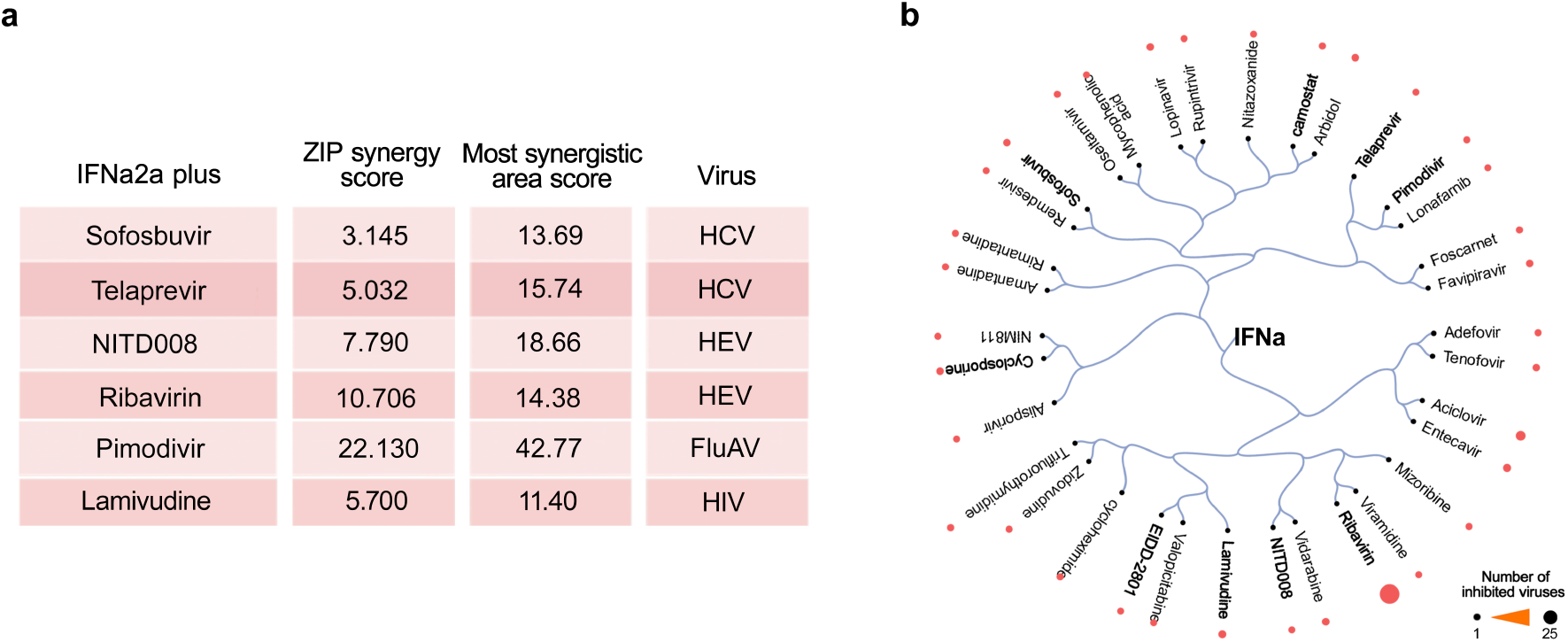
Synergistic IFNa-based combinations against other viruses. (**a**) Synergy and the most synergistic area scores of IFNa2a-based combinations against HCV, HEV, FluAV and HIV-1. (**b**) Structure-activity relationship of antivirals from known and novel (in bold) IFNa-based combinations. Compounds were extracted from IFNa-based combinations from https://antiviralcombi.info/ database. Compounds were clustered based on structural similarity calculated by ECPF4 fingerprints and visualized using the D3 JavaScript library. The broad-spectrum antiviral activities of the compounds are shown as bubbles, with larger bubbles corresponding to a larger number of targeted viruses.

Next, we studied IFNa2a in combination with known HEV inhibitors, NITD008 and ribavirin, against HEV infection in Huh-7.5 cells (Fig. S9, Fig. 8). Both NITD008 and ribavirin are nucleoside analogs which inhibit viral RNA synthesis. We observed that IFNa2a-NITD008 and IFNa2a-ribavirin were synergistic against HEV infection (ZIP synergy scores: 11 and 8; the most synergistic area scores: 14 and 19, respectively) while remaining nontoxic at synergistic doses for either drug.

We also tested IFNa2a in combination with known influenza inhibitor pimodivir against FluAV infection in A549 cells. Pimodivir (VX-787, JNJ-63623872) is an orally available anti-FluAV agent which targets viral polymerase basic protein 2, inhibits cap-snatching and has shown promising results in Phase II clinical trials [41, 42]. Cell viability was measured after 48 h in FluAV- and mock-infected cells to determine efficiency and toxicity of each compound and their combinations with IFNa2a (Fig. S9, Fig. 8). We observed that IFNa2a-pimodivir was synergistic against FluAV infection (ZIP synergy score: 22, the most synergistic area score: 43) while remaining nontoxic at synergistic doses for either drug.

Finally, we tested IFNa2a in combination with known anti-retroviral agent lamivudine against HIV-1 in TZM-bl cells. Lamivudine (3TC) is an orally available anti-HIV drug which inhibits viral reverse transcriptase [43]. Cell viability and HIV-induced luciferase expression were measured for each compound or their combination with IFNa2a after 48 h. We identified that treatment with IFNa2a and lamivudine was effective while being nontoxic at synergistic drug concentrations, with ZIP synergy scores of 6 and ZIP synergy score at the most synergistic area of 11 (Fig. S10, Fig. 8).

## Discussion

Currently, combinational therapies are still largely eschewed for the treatment of emerging viral infections in favor of monotherapies. This is in part due to the fact that many drug-drug interactions have not been fully explored or understood [44].

Here, we have reported several novel IFNa2a-based combination therapies that have better efficacy and lower toxicity than single drugs. In particular, we report novel *in vitro* activities of IFNa2a combinations with remdesivir, EIDD-2801, camostat, cycloheximide, and convalescent serum against SARS-CoV-2, with sofosbuvir or telaprevir against HCV infection, with NITD008 or ribavirin against HEV infection, with pimodivir against FluAV, as well as with lamivudine against HIV-1 infection. Our results indicate that other IFNa could be as efficient as IFNa2a when combined with these antivirals. Moreover, they expand the spectrum of antiviral activities of these combinations and emphasize the potential of IFNa-based combinatorial approach (Fig. 8b). Interestingly, pimodivir, lamivudine, remdesivir, EIDD-2801, NITD008, ribavirin and sofosbuvir interfere with synthesis of viral nucleic acids, whereas camostat, cycloheximide, telaprevir and convalescent serum inhibit other steps of viral replication cycle [33, 36, 41, 45], indicating that IFNa could be combined with virus- and host-directed agents targeting different stages of virus replication.

Based on our experiments, we propose the following mechanism of action of the IFNa-based combinations (Fig. S11). IFNa induces transcription of ISGs including IFIT1, IFIT2 and IFIT3, which recognize viral RNA; OASL and OAS2, which are involved in RNase L-mediated RNA degradation; and IDO1, which catalyzes kynurenine biosynthesis. IFNa also facilitates expression of several cytokines and virus-activated synthesis of IFNL1, IFNL2, IFNL3, IFNL4, and IFNB1, which alert the neighboring cells of upcoming infection. Therefore, combination treatments of IFNa and therapies targeting viral nucleic acid synthesis or other stages of virus replication can inhibit infection within a virus-host system.

Furthermore, we demonstrated anti-SARS-CoV-2 activity of IFNa-remdesivir in human lung organoids and hamsters. In particular, the combination treatment suppressed viral RNA expression more effectively than the drugs alone, while inducing transcription of antiviral genes. Thus, we have identified combination treatments that reduce viral replication at lower concentrations than is required with monotherapies. The low effective doses of these combination drugs may have several clinical advantages, including improved patient outcome and fewer adverse events.

In conclusion, the potential of using clinical grade IFNs as therapeutics against SARS-CoV-2 and other viral infections has raised interest recently [46]. It has been demonstrated that administration of IFNs in patients with early onset and mild symptoms inhibit infection and favor SARS-CoV-2 clearance [47, 48]. Our work may suggest that IFNa-based combinations with other antiviral agents may favor treatment of COVID-19 patients at various stages of disease and severity. It is conceivable that particularly vulnerable groups of COVID-19 patients with impaired immunity (i.e., impaired B-cell response, IFN response and T-cell response) may also benefit from combination of effective antivirals that are amplified by a dose of IFNa to elicit a clinical host response. We believe further development of IFNa-based combinations for treatment of SARS-CoV-2 and other viral infections can lead to practical therapeutic options that are more effective while having potentially reduced side effects than currently existing treatments. Moreover, the capacity to deliver IFNa-based combinations through different administration routes could allow for the treatment of patients at different stages of COVID-19 and other viral diseases [37, 49, 50].

## Materials and Methods

### Drugs, viruses, cells, lung organoids and hamsters

Table S1 lists IFNs and other antiviral agents, their suppliers and catalogue numbers. Lyophilized IFNs were dissolved in sterile deionized water to obtain 200 μg/mL concentrations. Compounds were dissolved in dimethyl sulfoxide (DMSO; Sigma-Aldrich, Hamburg, Germany) or milli-Q water to obtain 10 mM stock solutions. The convalescent serum (G614) from a recovered COVID-19 patient has been described in a previous study [25].

The propagation of wild-type SARS-CoV-2 (hCoV-19/Norway/Trondheim-S15/2020), recombinant mCherry-expressing SARS-CoV-2 strains (SARS-CoV-2-mCherry), wild type human influenza A/Udorn/307/1972 (H3N2), HCV and HIV-1 have been also described previously [25, 51–55]. SARS-CoV-2 strain Slovakia/SK-BMC5/2020 was provided by the European Virus Archive global (EVAg) and propagated in VeroE6/TMPRSS2 cells. To quantitate the production of infectious virions, we titered the viruses using plaque assays or ELISA [25, 51–54].

A plasmid harboring a sub-genomic HEV sequence coupled with a GFP reporter gene (Kernow-C1 p6 clone, gt3; GenBank Accession No. JQ679013) was used to generate HEV transcripts. Viral capped RNAs were transcribed *in vitro* from linearized plasmid using mMESSAGE mMACHINE^™^ T7 Transcription Kit (Thermofisher, USA). 1.5 x 10^7^ Huh-7.5 cells/mL in 400 μL of Maxcyte electroporation buffer were electroporated with 10 μg of p6-GFP sub-genomic HEV RNA. Electroporation was carried out with a Gene Pulser system (Bio-Rad, Munich, Germany) and allowed to recover for 30 min in a 37°C incubator. Recovered cells were resuspended in 10 mL prewarmed DMEM complete medium and maintained in an incubator for 24 h.

The propagation of human non-small cell lung cancer Calu-3; human adenocarcinoma alveolar basal epithelial A549; African green monkey kidney Vero-E6; T-cell like ACH-2 cells, which possess a single integrated copy of the provirus HIV-1 strain LAI (NIH AIDS Reagent Program); and human cervical cancer-derived TZM-bl, which express firefly luciferase under control of HIV-1 long terminal repeat (LTR) promoter allowing quantitation of the viral infection (tat-protein expression by integrated HIV-1 provirus) using firefly luciferase assay, have been described in our previous studies [25, 51–54]. Human hepatoma cells (Huh-7.5) were cultured in Dulbecco’s modified Eagle’s medium (DMEM) (Invitrogen, Karlsruhe, Germany) supplemented with 10% fetal bovine serum (Invitrogen), 1% nonessential amino acids (Invitrogen), 100 μg/mL of streptomycin (Invitrogen), and 100 IU/mL of penicillin.

The lung organoids (LOs) were generated as described previously (10.3390/v13040651). Briefly, induced pluripotent stem cells (IPSCs) were subjected to embryoid body induction using embryoid bodies (EB)/primitive streak media (10 μM Y-27632 and 3 ng/mL BMP4 in serum-free differentiation (SFD) media consisting of 375 mL Iscove’s Modified Dulbecco’s Medium (IMDM), 100 mL Ham’s F-12, 2.5 mL N2, 5 mL B27, 3,75 mL 7.5% BSA, 5 mL 1% penicillin–streptomycin, 5 mL GlutaMax, 50 μg/mL ascorbic acid, and 0.4 μM monothioglycerol) in ultra-low attachment plates. After 24 h the media was replaced with endoderm induction media (10 μM Y-27632, 0.5 ng/mL BMP4, 2.5 ng/mL FGF2, and 100 ng/mL Activin A in SFD media). Extra media was added every day for 3 days. The embryoid bodies were collected and dissociated using 0.05% Trypsin/EDTA and plated on fibronectin-coated plates with a cell density of 85,000 cells/cm^2^. Cells were then incubated in anteriorization media-1 (100 ng/mL Noggin, and 10 μM SB431542 in SFD media), followed by an incubation with anteriorization media-2 (10 μM SB431542, and 1 μM IWP2 in SFD media). The anteriorization media-2 was replaced with ventralization media (3 μM CHIR99021, 10 ng/mL FGF10, 10 ng/mL FGF7, 10 ng/mL BMP4, and 50 nM all-trans Retinoic acid in SFD media) and incubated for two days. The cell monolayer was then lifted by gentle pipetting, and the suspended cells were transferred to an ultra-low attachment plate where they would form the lung organoids.

Thirty-two 6-week-old healthy female Syrian hamsters were obtained from Janvier Labs. The animals were maintained in pathogen free health status according to the FELASA guidelines. The animals were individually identified and were maintained in housing rooms under controlled environmental conditions: temperature: 21 ± 2°C, humidity 55 ± 10%, photoperiod (12h light/12h dark), H14 filtered air, minimum of 12 air exchanges per hour with no recirculation. Each cage was labeled with a specific code. Animal enclosures provided sterile and adequate space with bedding material, food and water, environmental and social enrichment (group housing) as described below: IsoRat900N biocontainment system (Techniplast, France), poplar bedding (Select fine, Safe, France), A04 SP-10 diet (Safe, France), tap water, environmental enrichment, tunnel, wood sticks. Animal housing and experimental procedures were conducted according to the French and European Regulations and the National Research Council Guide for the Care and Use of Laboratory Animals. The animal BSL3 facility is authorized by the French authorities (Agreement N° D92-032-02). All animal procedures (including surgery, anesthesia, and euthanasia as applicable) were approved by the Institutional Animal Care and Use Committee of CEA and French authorities (CETEA DSV – n° 44).

### Drug Testing and Drug Sensitivity Quantification

Approximately 4 × 10^4^ Vero-E6 or Calu-3 cells were seeded per well in 96-well plates. The cells were grown for 24 h in DMEM or DMEM-F12, respectively, supplemented with 10% FBS and Pen– Strep. The medium was then replaced with DMEM or DMEM-F12 containing 0.2% BSA, Pen–Strep and the compounds in 3-fold dilutions at 7 different concentrations. No compounds were added to the control wells. The cells were infected with SARS-CoV-2 or SARS-CoV-2-mCherry strains at a moi of 0.01 or mock. After 72 or 48 h of infection, a CellTiter-Glo (CTG) assay was performed to measure cell viability. Drug efficacy on SARS-CoV-2-mCherry infected cells was measured on PFA- or acetone-fixed cells with fluorescence.

For testing compound toxicity and efficacy against FluAV, approximately 4 × 10^4^ A549 cells were seeded in each well of a 96-well plate. The cells were grown for 24 h in DMEM supplemented with 10% FBS and Pen–Strep. The medium was then replaced with DMEM containing 0.2% BSA, Pen–Strep, 0,5 μg/mL TPSK-trypsin and compounds in three-fold dilutions at seven different concentrations. No compounds were added to the control wells. The cells were infected with FluAV (moi = 0.5) or mock. At 48 hpi, the media was removed, and a CTG assay was performed to measure cell viability.

For testing compound toxicity and efficacy against HIV-1, approximately 4 × 10^4^ TZM-bl cells were seeded in each well of a 96-well plate in DMEM supplemented with 10% FBS and Pen–Strep. The cells were grown for 24 h in growth medium. The medium was then replaced with DMEM containing 0.2% BSA, Pen–Strep and the compounds in 3-fold dilutions at 7 different concentrations. No compounds were added to the control wells. The cells were infected with HIV-1 (corresponding to 300 ng/mL of HIV-1 p24) or mock. At 48 hours post-infection (hpi), the media was removed from the cells, the cells were lysed, and firefly luciferase activity was measured using the Luciferase Assay System (Promega, Madison, WI, USA). In a parallel experiment, a CTG assay was performed to measure cell viability.

We also examined cytotoxicity and antiviral activity of drug combinations using GFP-expressing HCV in Huh-7.5 cells by following previously described procedures [56]. For testing compound toxicity and efficacy against HEV, electroporated Huh-7.5 cells were seeded in the 384-well plate (3×10^3^ cells/well) with immune-modulators at indicated concentrations for 72 h. HEV replication was analyzed by determining the number of GFP-positive cells using fully automated confocal microscopy (Operetta CLS; PerkinElmer Devices).

The half-maximal cytotoxic concentration (CC_50_) for each compound was calculated based on viability/death curves obtained on mock-infected cells after non-linear regression analysis with a variable slope using GraphPad Prism software version 7.0a. The half-maximal effective concentrations (EC_50_) were calculated based on the analysis of the viability of infected cells by fitting drug dose-response curves using four-parameter (*4PL*) logistic function *f*(*x*):

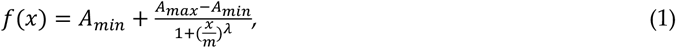

where *f*(*x*) is a response value at dose *x*, *A_min_* and *A_max_* are the upper and lower asymptotes (minimal and maximal drug effects), *m* is the dose that produces the half-maximal effect (EC_50_ or CC_50_), and *λ* is the steepness (slope) of the curve. The relative effectiveness of the drug was defined as selectivity index (*SI* = CC_50_/EC_50_).

To quantify each drug responses in a single metric, a drug sensitivity score (*DSS*) was calculated as a normalized version of standard area under dose-response curve (*AUC*), with the baseline noise subtracted, and normalized maximal response at the highest concentration (often corresponding to off-target toxicity):

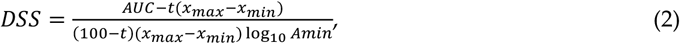

where activity threshold *t* equals 10%, and DSS ∈ [0,50].

### Drug Combination Testing and Synergy Calculations

Calu-3, A549, TZM-bl or Huh-7.5 cells were treated with different concentrations of two drugs and infected with SARS-CoV-2-mCherry (moi 0.01), FluAV (moi 0.5), HIV-1 (corresponding to 300 ng/mL of HIV-1 p24), HCV or mock. In addition, HEV-expressing Huh-7.5 cells were treated with different concentrations of two drugs. After 48 h, cell viability and reporter protein expression (SARS-CoV-2-mCherry, HIV-1, HCV-GFP, and HEV-GFP) were measured.

To test whether the drug combinations act synergistically, the observed responses were compared with expected combination responses. The expected responses were calculated based on the ZIP reference model using SynergyFinder version 2 [57, 58]. Final synergy scores were quantified as average excess response due to drug interactions (i.e., 10% of cell survival beyond the expected additivity between single drugs represents a synergy score of 10). Additionally, we calculated most synergistic area scores for each drug combination – the most synergistic 3-by-3 dose-window in dose-response matrixes.

LOs were treated with 0.5 μM remdesivir, 5 ng/mL IFNa2a, or their combination and infected with SARS-CoV-2-mCherry (moi 0.1). No compounds were added to the control wells. At 72 hpi, the LOs were stained using Cell Toxicity Green reagent (CTxG, Promega), and cell nuclei were stained with DAPI. Cells were fixed with PFA and imaged using microscopy. Representative images (*n* = 3) were selected.

### Prophylactic Study of Remdesivir, IFNaa and Their Combination Against SARS-CoV-2 Infection in Hamsters

Thirty-two animals were weighed and divided into 4 homogenous groups of 8 animals. Group 1 received 5 mL/kg vehicle IP 2h before infection. Group 2 received 40 μg/kg (5 mL/kg) mouse recombinant IFNa IP 2h before infection. Group 3 received 5mg/kg (5 mL/kg) remdesivir IP 2h before infection. Group 4 received a combination of 5mg/kg remdesivir and 40 μg/kg (5 mL/kg) IFNa IP 2h before infection. All groups received SARS-CoV-2 intranasally. Animal viability, behavior and clinical parameters were monitored daily. After 3 days animals were deeply anesthetized using a cocktail of 30 mg/kg (0.6 mL/kg) Zoletil and 10 mg/kg (0.5 mL/kg) Xylazine IP. Cervical dislocation followed by thoracotomy was performed before lung collection. The entire left lungs and superior, middle, post-caval and inferior lobes of right lungs were put in RNAlater tissue storage reagent overnight at 4°C, then stored at −80°C until RNA extraction.

### RT-qPCR Analysis

Viral RNA was extracted using the QIAamp Viral RNA Mini Kit (Qiagen). RT-PCR was performed using SuperScript^™^ III One-Step qRT-PCR System kit (commercial kit #1732-020, Life Technologies) with primers nCoV_IP2-12669Fw: ATGAGCTTAGTCCTGTTG, nCoV_IP2-12759Rv: CTCCCTTTGTTGTGTTGT, and nCoV_IP2-12696bProbe(+): Hex-AGATGTCTTGTGCTGCCGGTA-BHQ-1 or nCoV_IP4-14059Fw: GGTAACTGGTATGATTTCG, nCoV_IP4-14146Rv: CTGGTCAAGGTTAATATAG, and nCoV_IP4-14084Probe(+): Fam-TCATACAAACCACGCCAGG-BHQ-1 targeting IP2 and IP4 regions SARS-CoV-2 RdRP gene as well as ORF1ab_Fw: CCGCAAGGTTCTTCTTCGTAAG, ORF1ab_Rv: TGCTATGTTTAGTGTTCCAGTTTTC, ORF1ab_probe: Hex-AAGGATCAGTGCCAAGCTCGTCGCC-BHQ-1 targeting another region on ORF1ab. RT-qPCR was performed using a Bio-Rad CFX384^™^and adjoining software. The relative gene expression differences were calculated using β-Actin as control and the results were represented as relative units (RU). Technical triplicates of each sample were performed on the same qPCR plate and non-templates and non-reverse transcriptase samples were analysed as negative controls. Statistical significance (p < 0.05) of the quantitation results was evaluated with t-test. Benjamini-Hochberg method was used to adjust the p-values.

### Gene Expression Analysis

Calu-3 cells, LOs or Syrian hamsters were treated with drugs or vehicles and infected with SARS-CoV-2, SARS-CoV-2-mCherry or mock. Total RNA was isolated using RNeasy Plus Mini kit (Qiagen, Hilden, Germany) from Calu-3 cells, LOs or lungs of Syrian hamsters. Polyadenylated mRNA was isolated from 250 ng of total RNA with NEBNext Poly(A) mRNA magnetic isolation module. NEBNext Ultra II Directional RNA Library Prep kit from Illumina was used to prepare samples for sequencing. Sequencing was done on NextSeq 500 instrument (set up: single-end 1 x 76 bp + dual index 8 bp) using NextSeq High Output 75 cycle sequencing kit (up to 400M reads per flow cell). Reads were aligned using the Bowtie 2 software package version 2.4.2 to the NCBI reference sequence for SARS-CoV-2 (NC_045512.2) and to the human GRCh38 genome. The number of mapped and unmapped reads that aligned to each gene were obtained with the featureCounts function from Rsubread R-package version 2.10. The GTF table for the SARS-CoV-2 reference sequence was downloaded from https://ftp.ncbi.nlm.nih.gov/genomes/all/GCF/009/858/895/GCF_009858895.2_SM985889v3/GCF_009858895.2_ASM985889v3_genomic.gtf.gz. The heatmaps were generated using the pheatmap package (https://cran.r-project.org/web/packages/pheatmap/index.html) based on log2-transformed or non-transformed profiling data.

### Metabolic Analysis

Calu-3 cells or LOs were treated with drugs or vehicle and infected with SARS-CoV-2, SARS-CoV-2-mCherry or mock. Metabolites were extracted from Calu-3 cells and LOs supernatants or from lung extracts. 100 μL of culture medium/lung extracts were mixed with 400 μL of cold extraction solvent (acetonitrile:methanol:water 40:40:20). Subsequently, samples were sonicated for 3 cycles (60 s, power = 60 and frequency = 37), vortexed for 2 min and centrifuged at 4 °C, 15000 g for 10 min. The supernatant was transferred to autosampler vials for LC-MS analysis. The extracts were analyzed with Thermo Vanquish UHPLC+ system coupled to a QExactive Orbitrap quadrupole mass spectrometer equipped with a heated electrospray ionization (H-ESI) source probe (Thermo Fischer Scientific, Waltham, MA, USA). A SeQuant ZIC-pHILIC (2.1 × 100 mm, 5 μm particles) HILIC phase analytical column (Merck KGaA, Darmstadt, Germany) was used as a chromatographic separation column.

Gradient elution was carried out with a flow rate of 0.1 mL/min with 20 mM ammonium carbonate, adjusted to pH 9.4 with ammonium solution (25%) as mobile phase A and acetonitrile as mobile phase B. The gradient elution was initiated from 20% mobile phase A and 80% mobile phase B and maintained for 2 min. Then, mobile phase A was gradually increased up to 80% for 17 min, followed by a decrease to 20% over the course of 17.1 min. and sustained for up to 24 min.

The column oven and auto-sampler temperatures were set to 40 ± 3 °C and 5 ± 3 °C, respectively. The mass spectrometer was equipped with a heated electrospray ionization (H-ESI) source using polarity switching and the following settings: resolution of 35,000, the spray voltages of 4250 V for positive and 3250 V for negative mode, sheath gas at 25 arbitrary units (AU), the auxiliary gas at 15 AU, sweep gas flow of 0, Capillary temperature of 275°C, and S-lens RF level of 50.0. Instrument control was operated with Xcalibur 4.1.31.9 software (Thermo Fischer Scientific, Waltham, MA, USA). Metabolite peaks were confirmed using the mass spectrometry metabolite library kit MSMLS-1EA (Sigma Aldrich supplied by IROA Technologies).

For data processing, final peak integration was done with the TraceFinder 4.1 software (Thermo Fisher Scientific, Waltham, MA, USA) and for further data analysis, the peak area data was exported as an Excel file. Data quality was monitored throughout the run using pooled healthy human serum as Quality Control (QC), which was processed and extracted in the same manner as unknown samples. After integration of QC data with TraceFinder 4.1, each detected metabolite was checked and %RSD were calculated, while the acceptance limit was set to ≤ 20%.

Blank samples were injected after every five runs to monitor any metabolite carryover. A carryover limit of ≤ 20% was set for each metabolite. Percentage background noise was calculated by injecting a blank sample at the beginning of the run. The acceptance limit for background noise was set at ≤ 20% for each metabolite.

## Ethics approval and consent to participate

Standard operational procedures were approved by institutional safety committee.

## Consent for publication

All authors have read and agreed to the published version of the manuscript.

## Availability of data and material

All data generated or analyzed during this study are included in this published article and its supplementary information files.

## Author Contributions

All authors contributed to the methodology, software, validation, formal analysis, investigation, resources, data curation, writing, and review and editing of the manuscript. D.K. conceptualized, supervised, and administrated the study and acquired funding.

## Acknowledgments

We thank personnel of Biomedicum function genomics (FuGu), Juho Vaananen and Martyn unit for transcriptomics analysis.

