## Supplementary table and figures for "Interferon alpha-based combinations suppress SARS-CoV-2 infection in vitro and in vivo"

**Supplementary data**

Table S1. Compounds, their suppliers, and catalogue numbers.

| **Name** | Cat # | **Company** |
| --- | --- | --- |
| Recombinant human IFNa1b | 11343594 | ImmunoTools |
| Recombinant human IFNa2a | 11343504 | ImmunoTools |
| Recombinant human IFNa2b | 11343514 | ImmunoTools |
| Recombinant human IFNb1a | 11343520 | ImmunoTools |
| Recombinant human IFNb1b | 11343542 | ImmunoTools |
| Recombinant human IFNg | 11343534 | ImmunoTools |
| Recombinant human IFNw1 | 11344784 | ImmunoTools |
| Recombinant human IL28A | 11340280 | ImmunoTools |
| Recombinant human IL‐29 | 11340290 | ImmunoTools |
| Recombinant mouse IFNaa | 10150-IF-050 | R&D Systems |
| Camostat mesylate | 16018 | Cayman Chemicals |
| Remdesivir | 30354 | Cayman Chemicals |
| Lamivudine | S1706 | Selleckchem |
| Cycloheximide | C7698-1g | SigmaAldrich |
| Pimodivir | HY-12353A/CS | MedChemExpress |
| EIDD-2801 | HY-135853 | MedChemExpress |
| Ribavirin | 0219606650 | MP Biomedicals |
| NITD008 | SML2409 | SigmaAldrich |
| Sofosbuvir | S2794 | Selleckchem |
| Telaprevir | S1538 | Selleckchem |

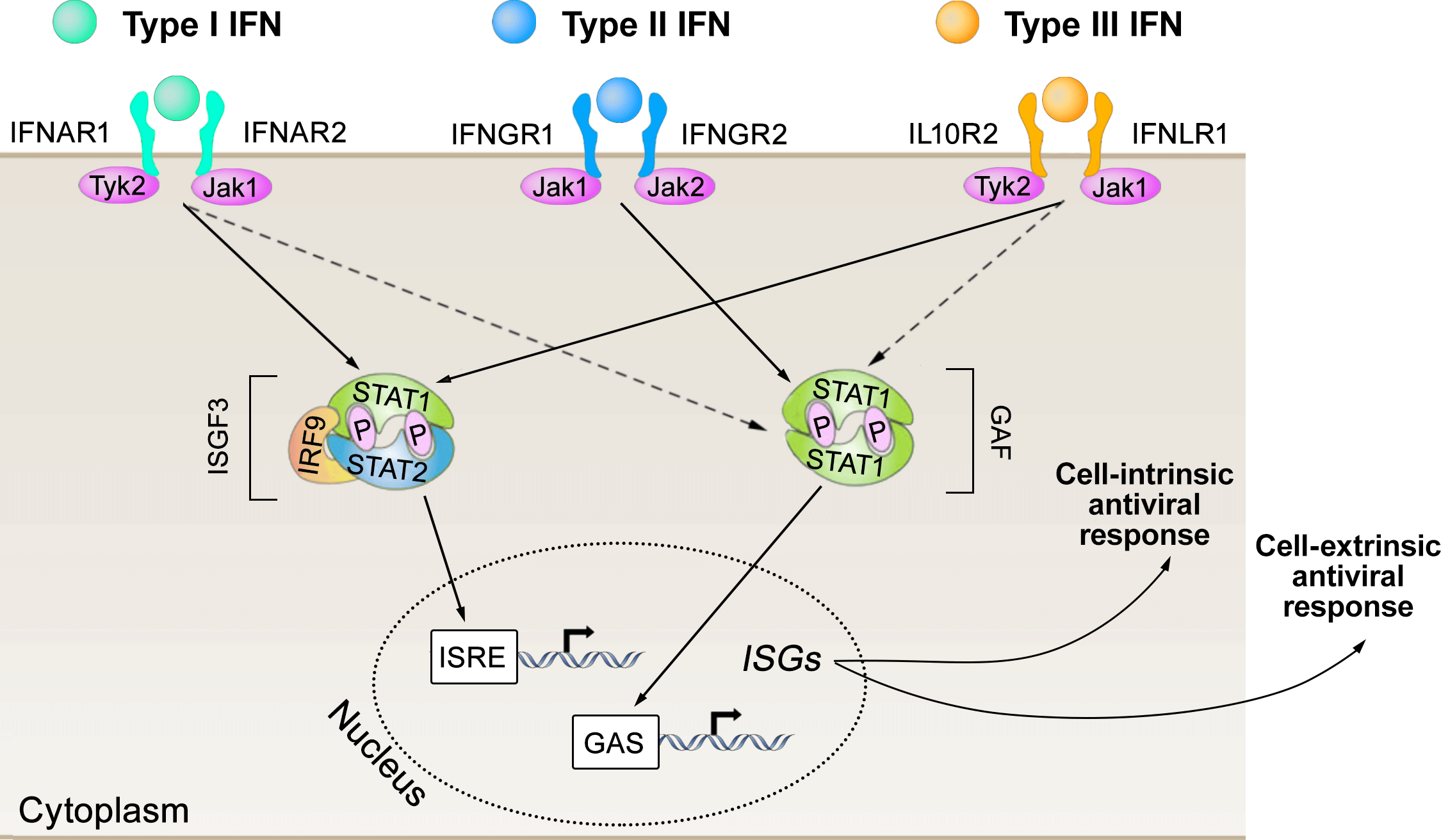

**Fig. S1.** Three types of IFNs and their targets (adapted from ^10^).

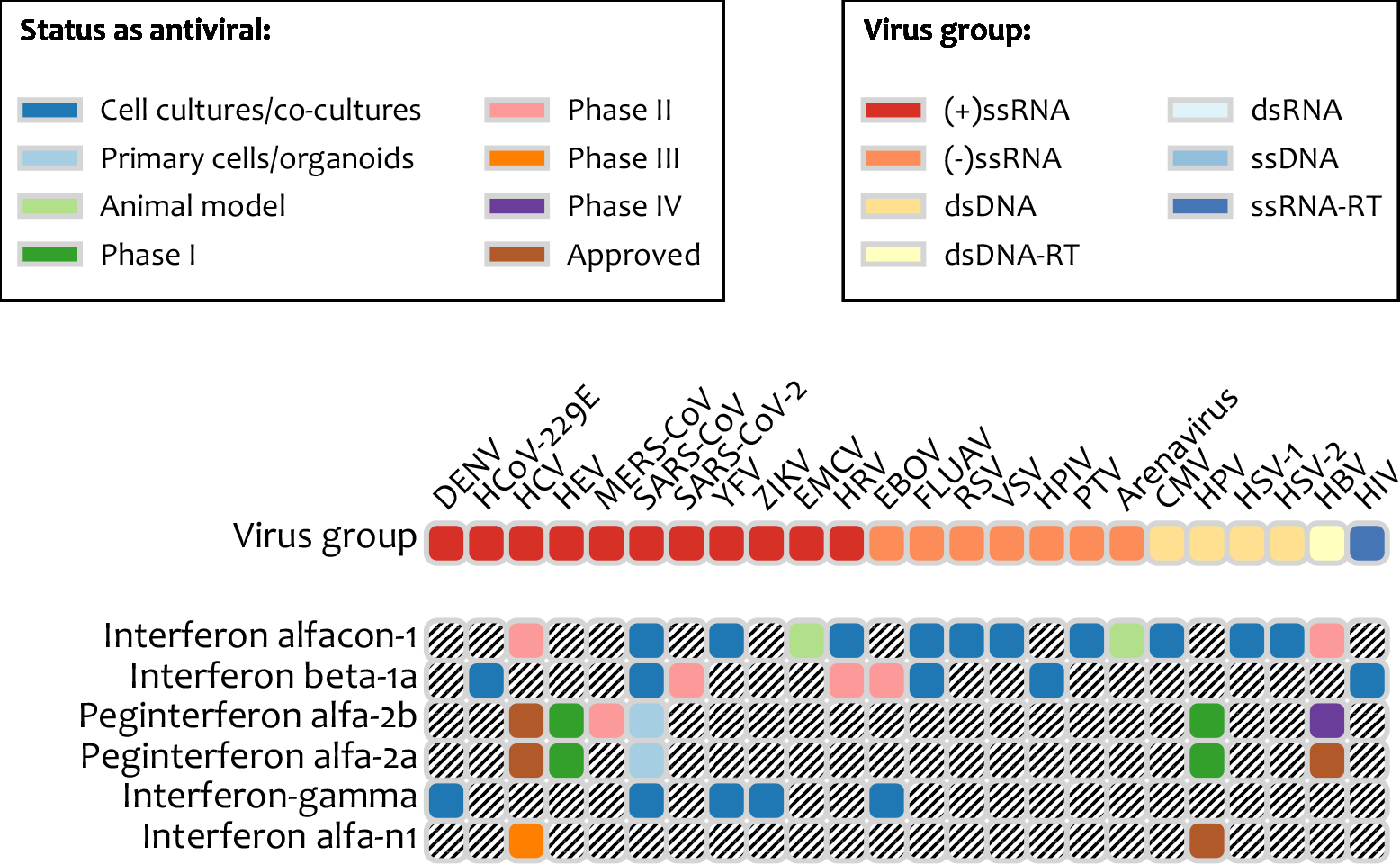

**Fig. S2.** Developmental statuses of several natural and recombinant human IFNs against a range of pathogenic human viruses. Data was retrieved from our database of broad-spectrum antivirals (<http://drugvirus.info/>).

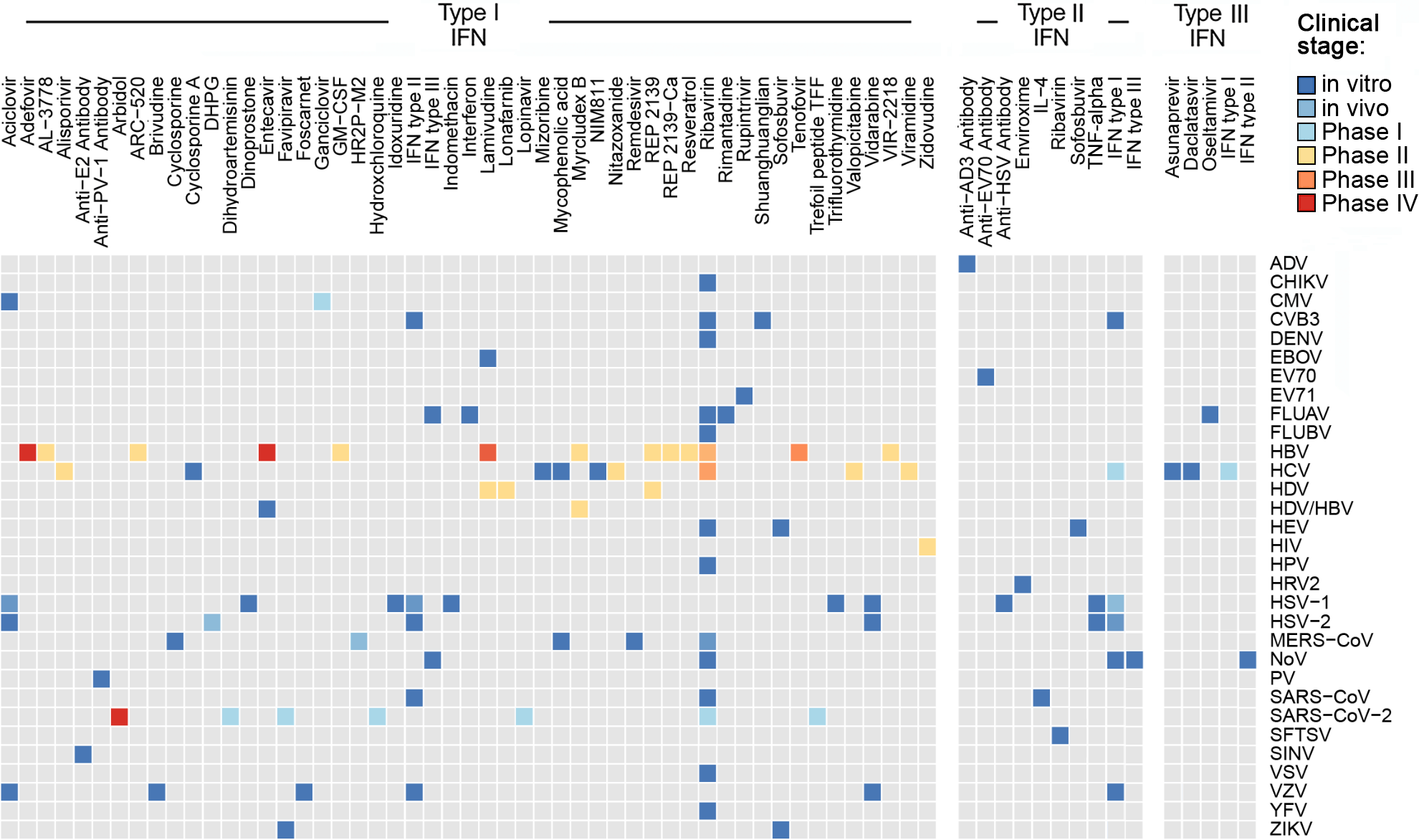

**Fig. S3.** Examples of IFN-based combinations and their developmental statuses. Data was retrieved from our antiviral drug combinations database ( https://antiviralcombi.info).

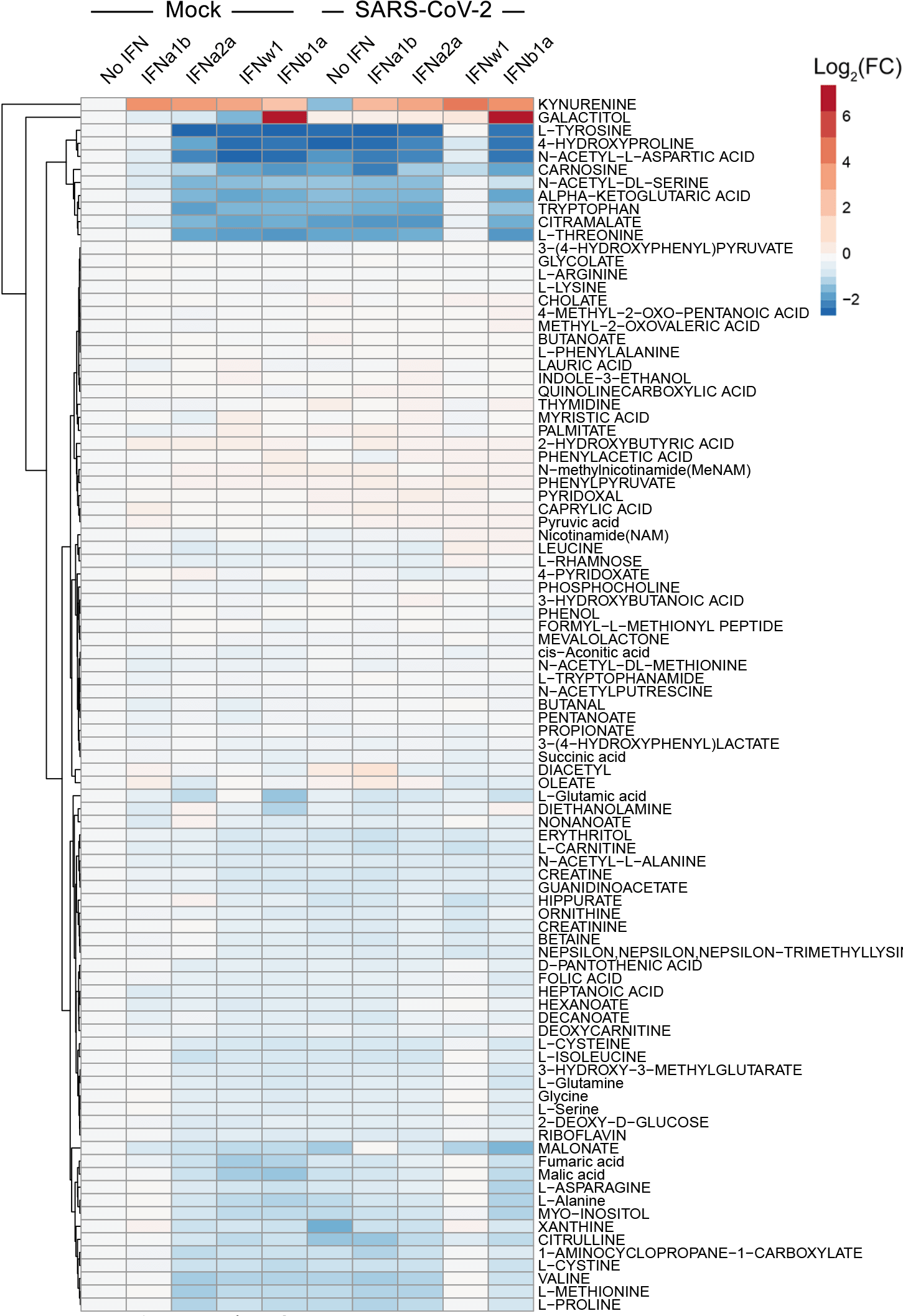

**Fig. S4**. Metabolomic profiles of mock- and SARS-CoV-2-infected Calu-3 cells non-treated or treated with type I IFNs.

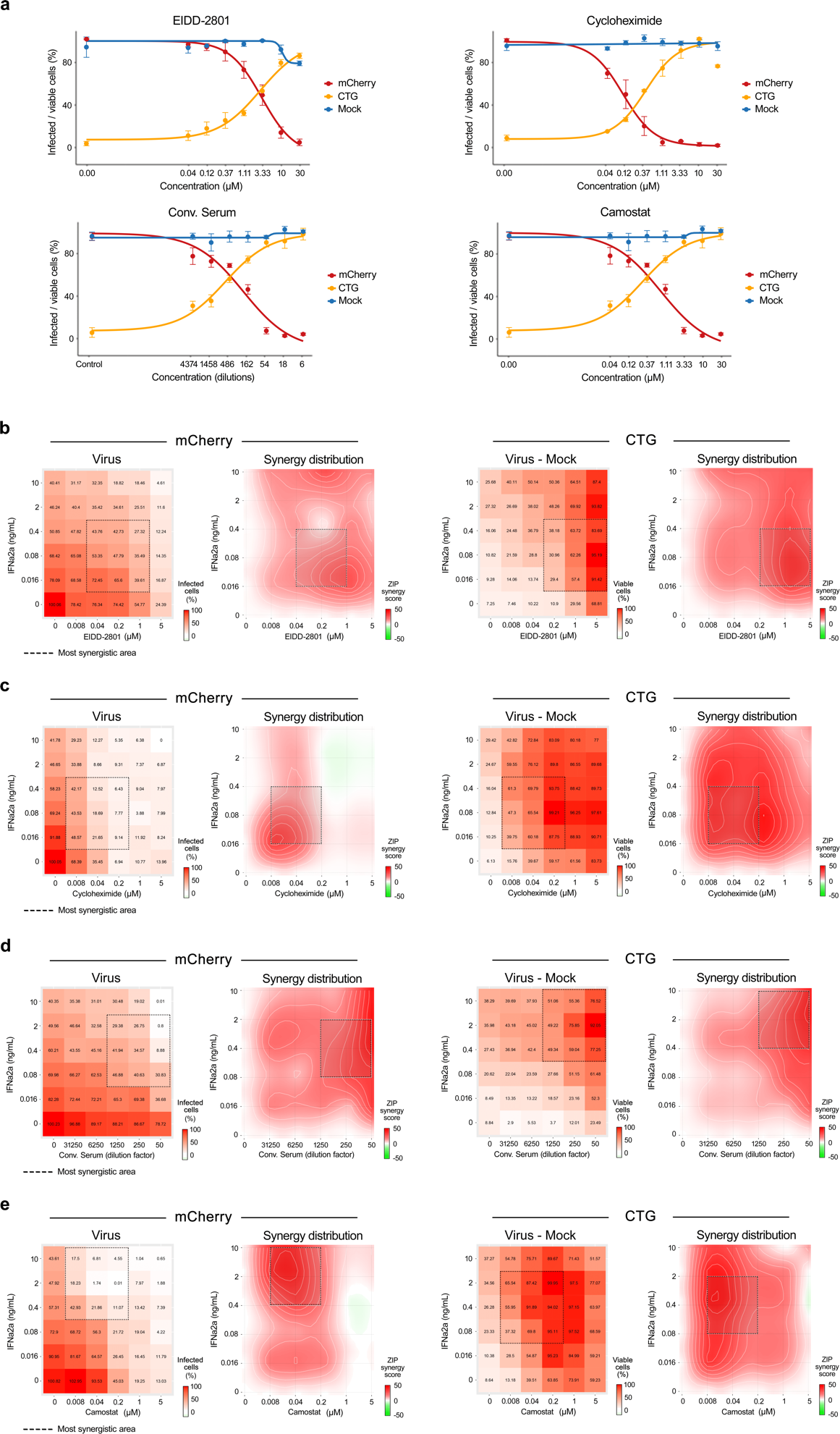

**Fig. S5**. Synergistic IFNa2a-based combinations against SARS-CoV-2-mCherry infection in Calu-3 cells. (**a**) Calu-3 cells were treated with increasing concentrations of a drug and infected with the SARS-CoV-2-mCherry or mock. After 48 h, the virus-mediated mCherry expression was measured (red curves). After 72 h, viability of virus- and mock-infected cells was determined using a CTG assay (yellow and blue curves, respectively). Mean ± SD; n = 3. (**b-e**, left panels) The 6 × 6 dose–response matrices and interaction landscapes of IFNa2a and remdesivir obtained using fluorescence analysis of SARS-CoV-2-mCherry-infected Calu-3 cells. ZIP synergy score was calculated for the drug combinations. (**b-e**, right panels) The 6 × 6 dose–response matrices and interaction landscapes of IFNa2a and remdesivir obtained using a cell viability assay (CTG) on mock-, and SARS-CoV-2-mCherry-infected Calu-3 cells. The selectivity for the indicated drug concentrations was calculated (selectivity = efficacy-(100-Toxicity)). ZIP synergy scores were calculated for indicated drug combinations.

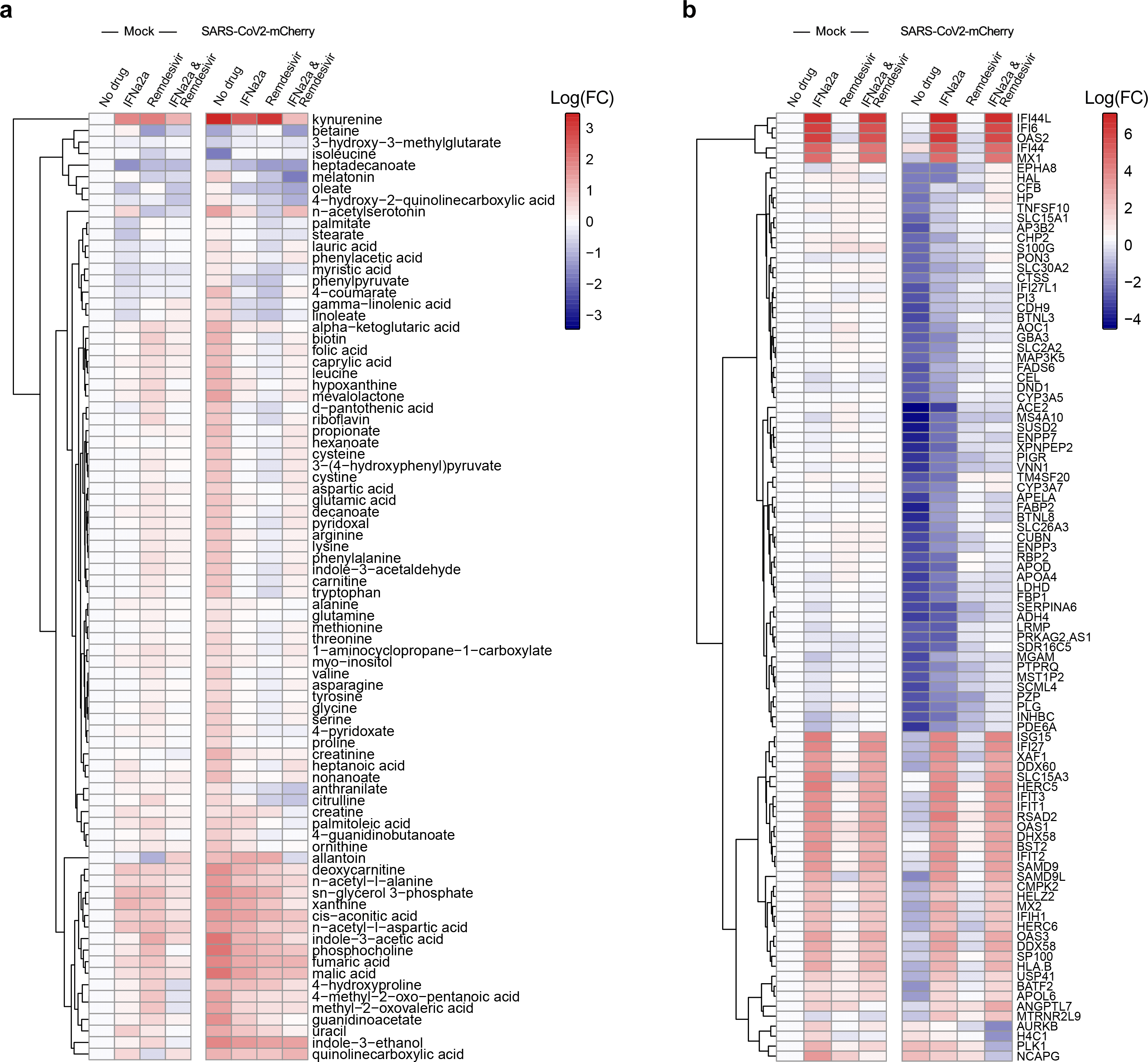

**Fig. S6.** Effect of IFNa2a-remdesivir on transcription of host genes in human lung organoids (LOs). LOs were treated with 0,5 μM remdesivir, 5 ng/mL IFNa2a, a combination thereof, or vehicle; then infected with SARS-CoV-2-mCherry (moi = 0,1) or mock. After 48 h, total RNA was extracted and sequenced. A heatmap of the most variable cellular genes affected by treatment and virus infection is shown. Each cell is colored according to the log2‐transformed expression values of the samples, expressed as fold‐change relative to the nontreated mock-infected control. Cut-off - 2.

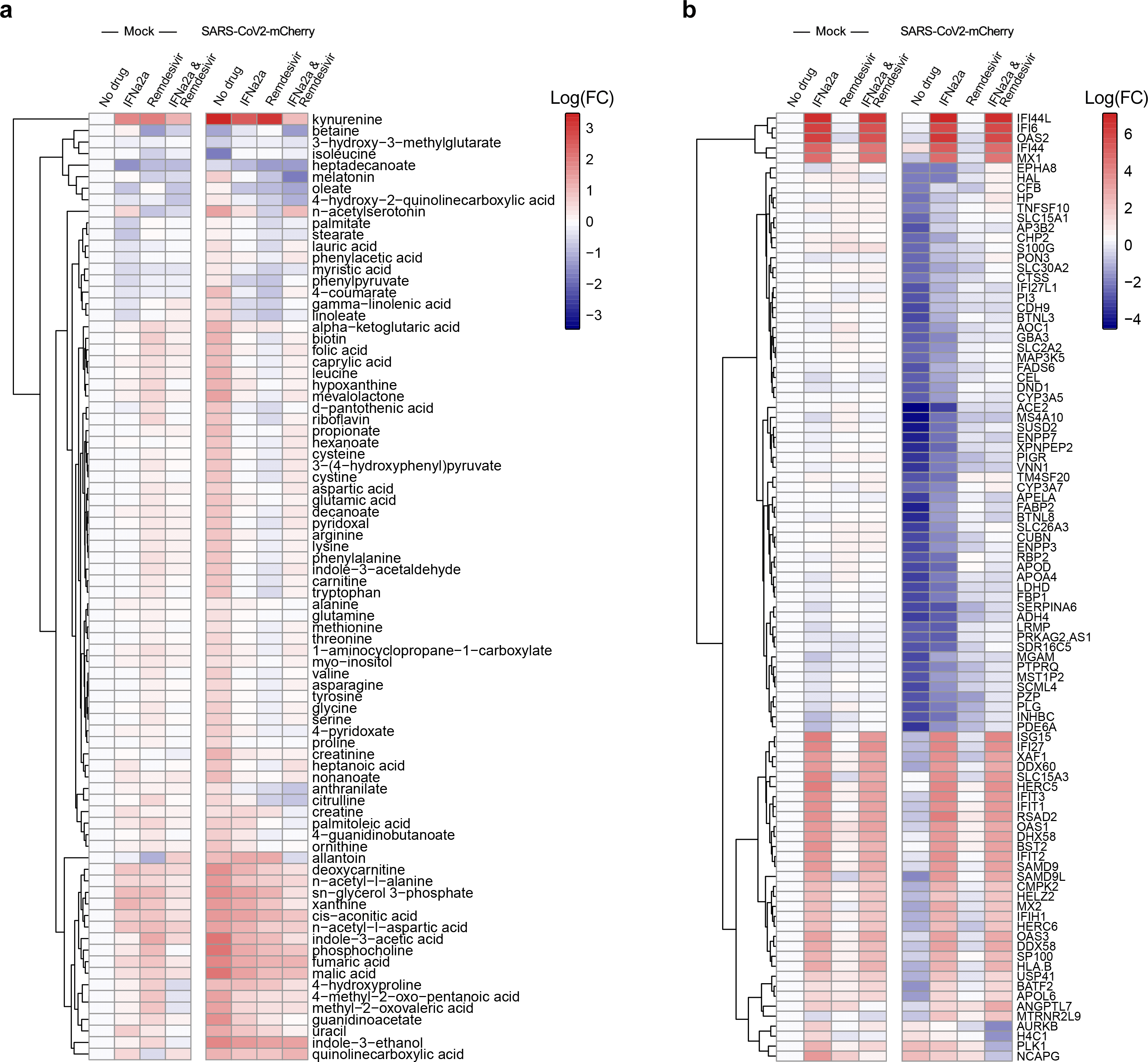

**Fig. S7.** Effect of IFNa2a-remdesivir on metabolism in human lung organoids (LOs). LOs were treated with 0,5 μM remdesivir, 5 ng/mL IFNa2a, a combination thereof, or vehicle; then infected with SARS-CoV-2-mCherry (moi = 0,1) or mock. After 48 h, the cell culture supernatants were collected, and metabolite levels were determined by LC‐MS/MS. A heatmap of the most affected metabolites is shown. Each cell is colored according to the log2‐transferred profiling values of samples, expressed as fold‐change relative to the mock control.

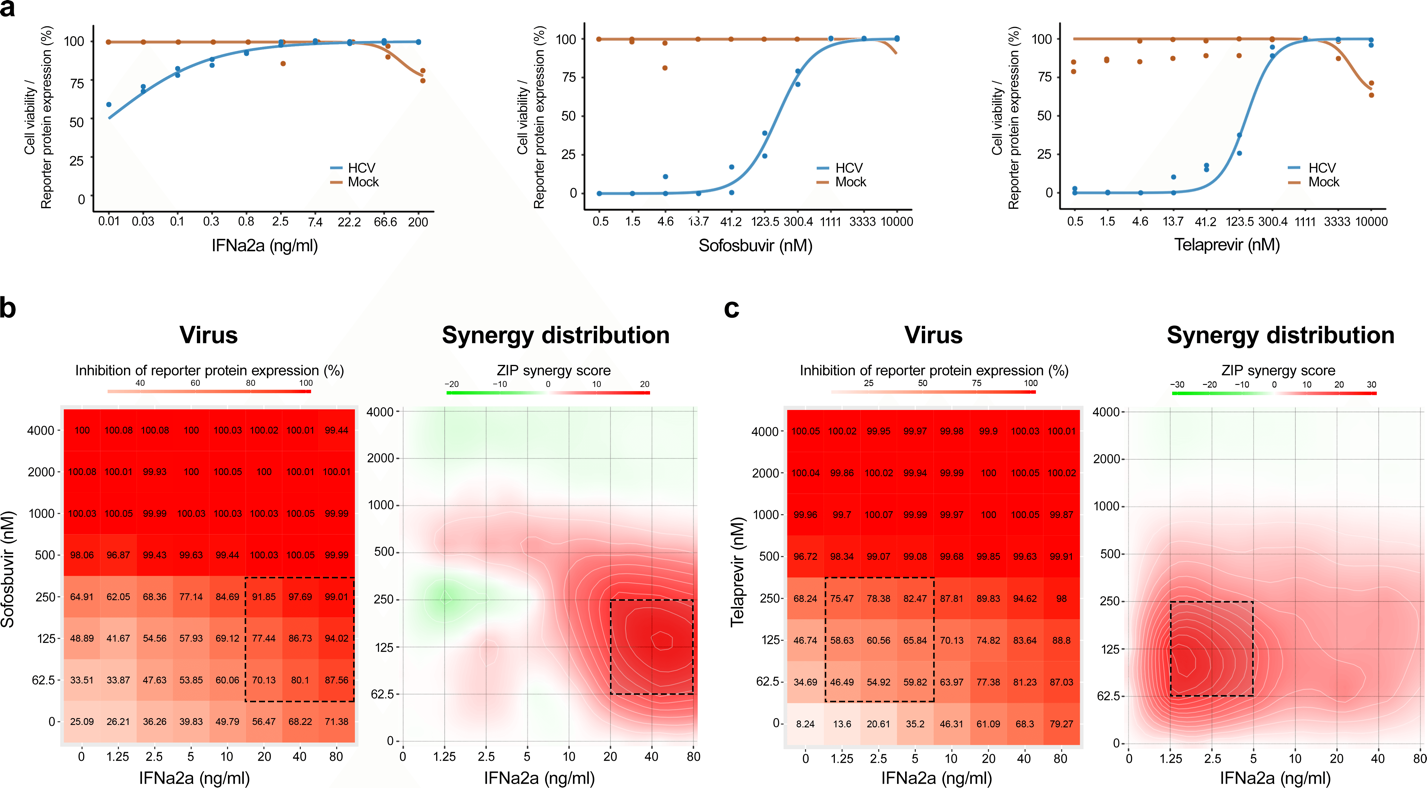

**Fig. S8.** Combinations of IFNa2a with sofosbuvir or telaprevir reduce HCV-mediated GFP expression in Huh-7.5 cells. (**a**) Huh-7.5 cells were treated with increasing concentrations of IFNa2a, sofosbuvir, or telaprevir and infected with HCV. After 72 h, the HCV-mediated GFP expression was measured (blue curves). The total number of cells was used as marker for cytotoxicity (red curves). Mean ± SD; n = 3. (**b**,**c**) The interaction landscape of IFNa2a-sofosbuvir and IFNa2a-telaprevir, measured using HCV-mediated GFP expression in Huh-7.5 cells (left panels). The interaction landscapes of drug combinations (right panels).

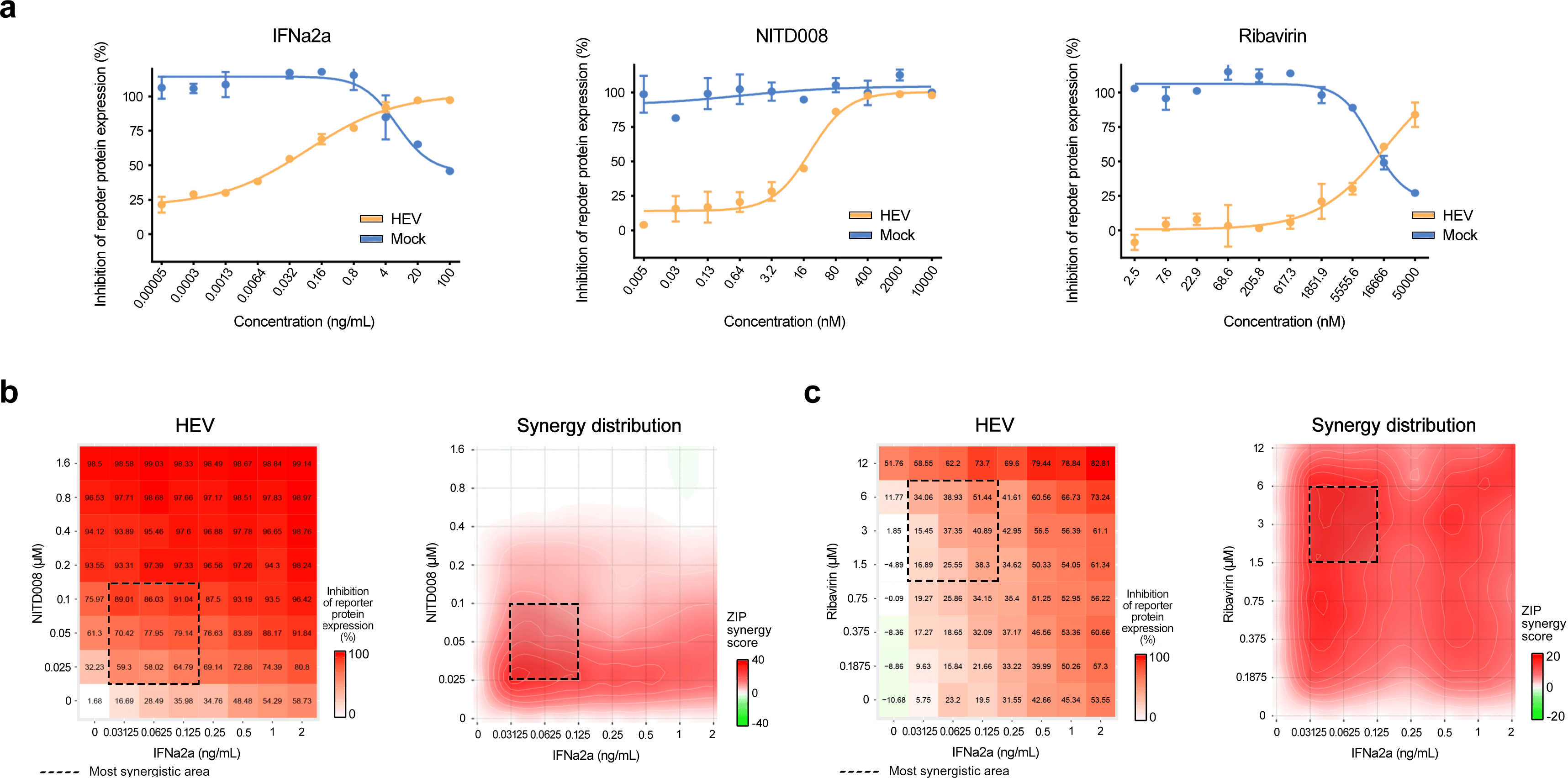

**Fig. S9.** Combinations of IFNa2a with ribavirin or NITD008 reduce HEV-mediated GFP expression in Huh-7.5 cells. (**a**) Huh-7.5 cells were transfected with p6-GFP sub-genomic HEV RNA or mock. After 24 h cells were treated with increasing concentrations of IFNa2a, NITD008 or ribavirin. After 72 h, the HEV-mediated GFP expression was measured (blue curves). Total number of cells was used as marker for cytotoxicity (red curves). Mean ± SD; n = 3. (**b**,**c**) The interaction landscape of IFNa2a-ribavirin and IFNa2a-NITDD008 were measured using HEV-mediated GFP expression in Huh-7.5 cells (left panels). The interaction landscapes of drug combinations (right panels).

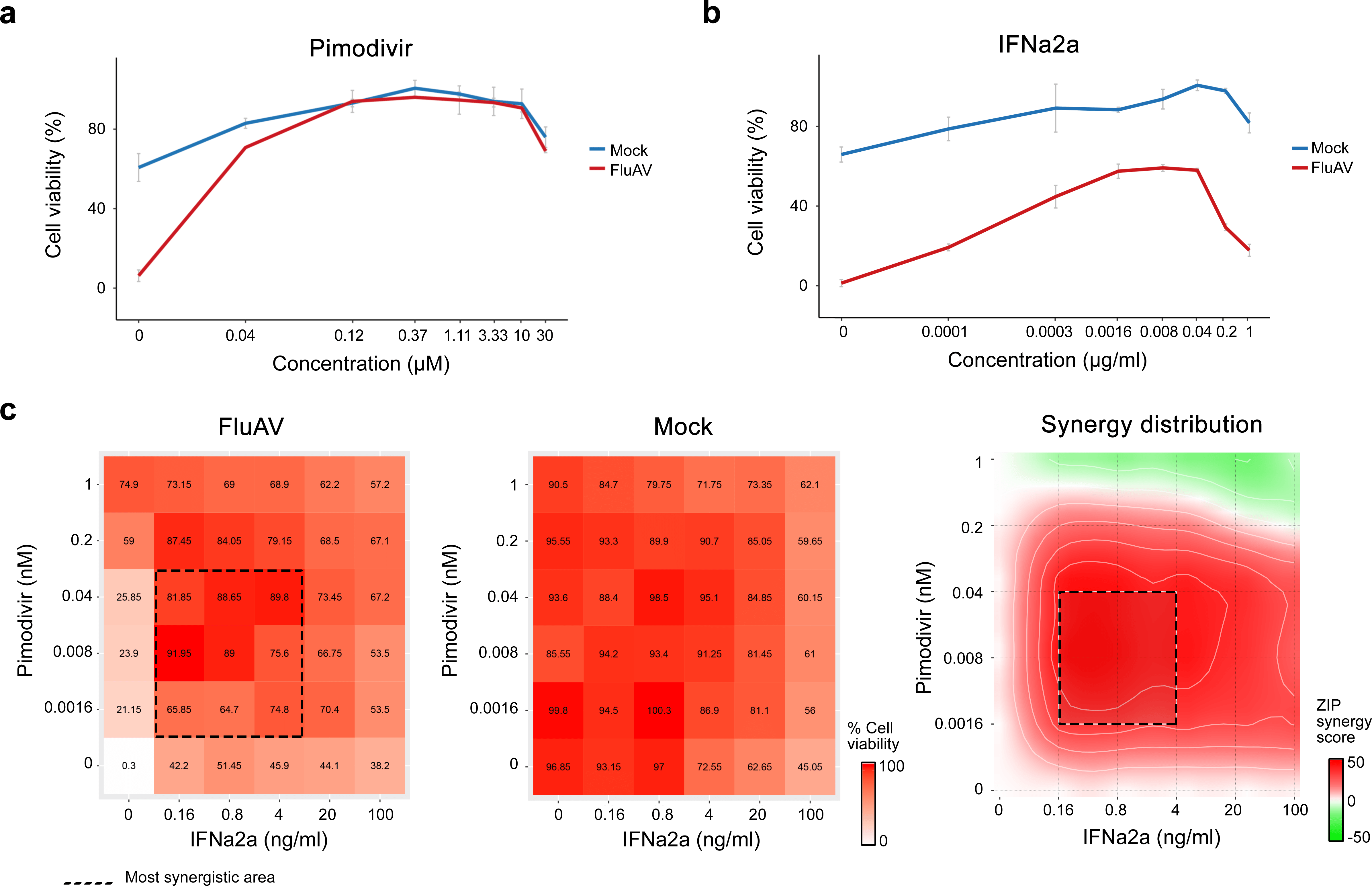

**Fig. S10.** Combination of pimodivir-IFNa2a reduces FluAV infection in A549 cells. (**a**,**b**) A549 cells were treated with increasing concentrations of pimodivir or IFNa2a and infected with the FluAV (moi = 0.5) or mock. After 48 h, cell viability was determined using a CTG assay. Mean ± SD; n = 3. (**c**) The interaction landscape of IFNa2a and pimodivir in FluAV- and mock infected A549 cells measured using CTG (left panels). The interaction landscape of both drugs showing synergy of the drug combination (right panel).

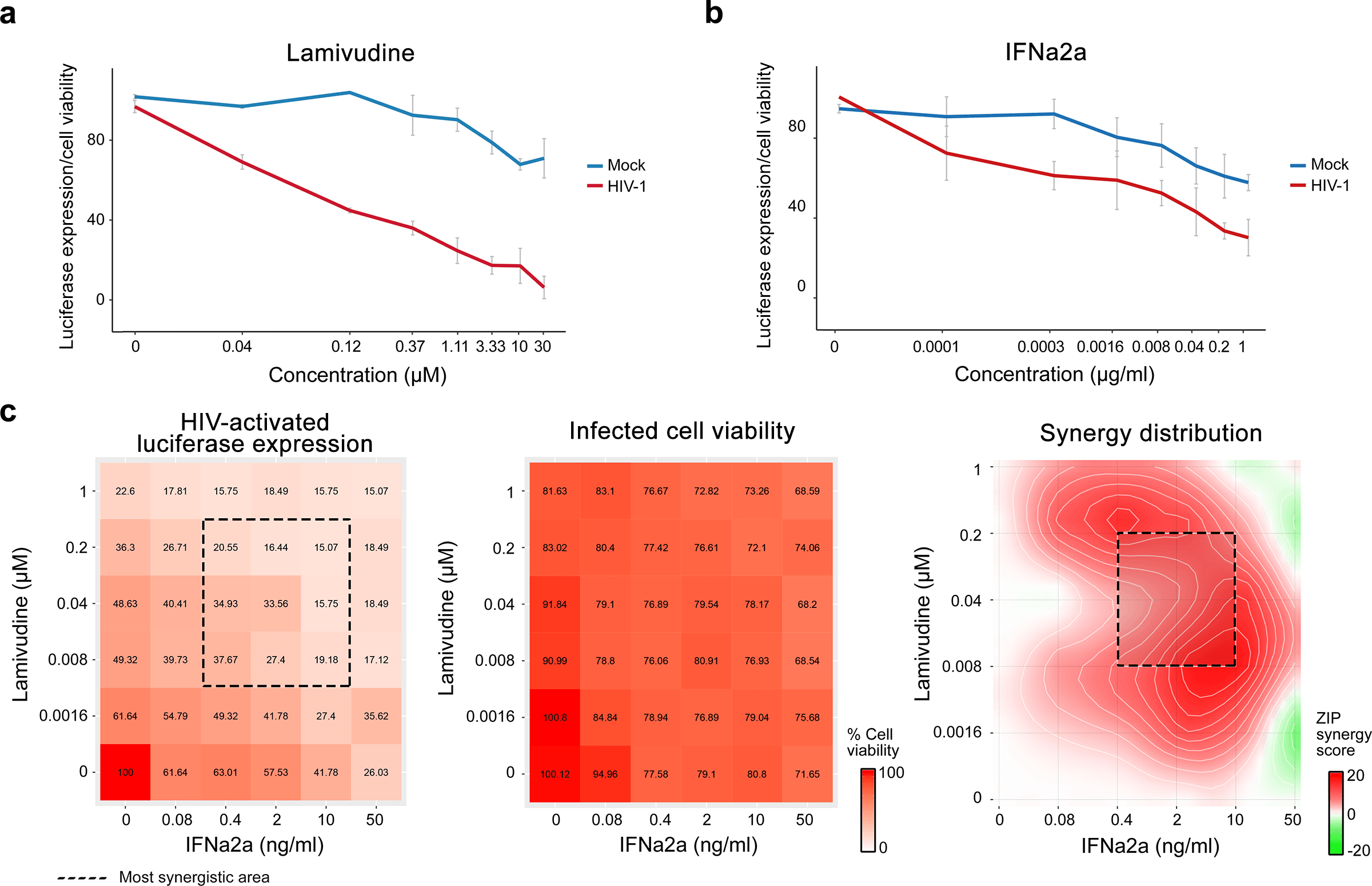

**Fig. S11.** Combination of lamivudine-IFNa2a reduces HIV-1 infection in TZM-bl cells. (**a**,**b**) TZM-bl cells were treated with increasing concentrations of lamivudine or IFNa2a and infected with the HIV-1 or mock. After 48 h, the HIV-activated luciferase expression was measured (red curves). Viability of mock-infected cells was determined using the CTG assay (blue curves). Mean ± SD; n = 3. (**c**) The interaction landscape of IFNa2a and lamivudine measured using HIV-1-activated luciferase expression in virus- and mock-infected TZM-bl cells, respectively (left panels). The interaction landscape of both drugs showing synergy of their combination (right panel).

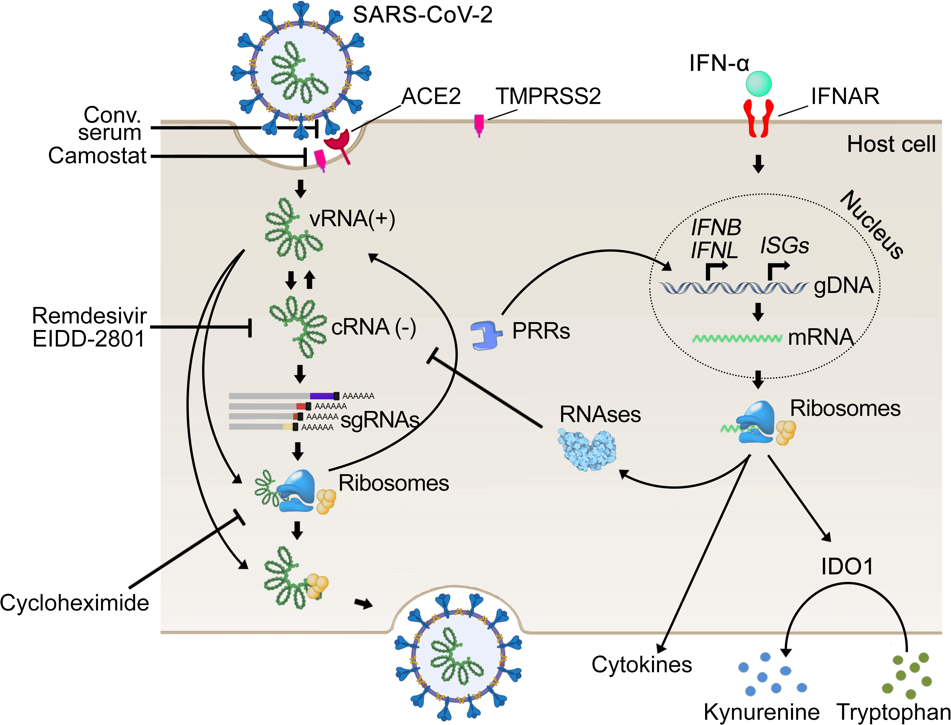

**Fig. S12**. Schematic representation of mechanisms of anti-SARS-CoV-2 actions of remdesivir, EIDD-2801, camostat, cycloheximide, and convalescent serum, and stages of virus replication cycle they target.
